# Electrophysiological and neuroimmune responses of a cortical tissue model to chronic gamma radiation exposure

**DOI:** 10.64898/2026.09.16.752167

**Authors:** Neal M. Lojek, Nazli S. Bostanci, Paulo Henrique Borges, Steven Snay, Andrew M. Rogers, Erno Sajo, Egle Cekanaviciute, Bryan J. Black, Chiara E. Ghezzi

**Affiliations:** Department of Biomedical Engineering, University of Massachusetts Lowell, Lowell MA, 01854, MA, USA; Department of Physics and Applied Physics, University of Massachusetts Lowell, Lowell MA, 01854, MA, USA; Space Biosciences Division, NASA Ames Research Center, Moffett Field, CA 94035, USA

**Keywords:** space radiation, electrophysiology, 3D tissue model, radioprotectant

## Abstract

Exposure to chronic space radiation is a major health concern for long-duration missions beyond low Earth orbit. While most experimental studies simulate mission-relevant doses using acute irradiation, astronauts will experience persistent low-dose-rate exposure over months. Here, we used engineered three-dimensional (3D) mouse cortical tissue models and two-dimensional (2D) primary cortical cultures to investigate the effects of radiation dose rate on neuroglial function, inflammatory responses, and neuronal network activity. Cultures were exposed to a cumulative 0.5 Gy dose of γ-radiation delivered either acutely (1 h) or chronically (166 h), with or without pretreatment using the radioprotective agent amifostine. In 3D cortical tissue models, we quantified DNA damage, astrocyte and microglia reactivity, neuronal survival, neurite morphology, and secretion of pro-inflammatory cytokines. In parallel, microelectrode array recordings were used to assess electrophysiological function in 2D neuronal networks following γ-radiation or treatment with the radiomimetic drug bleomycin. Acute gamma radiation induced modest astrocyte activation, whereas chronic exposure caused limited neuroimmune alterations. Neither exposure paradigm impaired neuronal network activity, despite measurable DNA damage responses. Together, these findings indicate that neuronal function is preserved following low-dose-rate gamma irradiation while glial populations display selective sensitivity to radiation exposure.

## Introduction

The detrimental central nervous system (CNS) effects of prolonged space radiation exposure are a major concern for crewed missions beyond low earth orbit, including lunar and Mars explorations. The health effects of space radiation on the CNS are expected to be of high likelihood and high consequence and, as such, require mitigation for mission success (Patel, Brunstetter et al. 2020). Human data on space radiation exposure is highly limited and estimates of clinical outcomes are derived from ground-based experiments utilizing ionizing radiation (IR) exposure of animal and tissue culture models to examine injury mechanisms (Huff, Poignant et al. 2023). Animal studies which use ground-based simulated space radiation indicate that behavioral changes and neurocognitive deficits are a concern (Alaghband, Klein et al. 2023, Ma, Li et al. 2024). However, these models are limited, as neurobehavioral readouts are often subjective (Schielke, Hartel et al. 2020) and most radiation exposure studies are treated with acute rather than chronic radiation. Additionally, whole animal models do not enable isolation and subsequent fundamental understanding of CNS-specific effects and those due to whole body exposure. While whole body effects are undoubtedly relevant to astronaut health, isolating the effects of IR on the CNS may be pivotal for developing our fundamental understanding of cell-type specific vulnerability to injury as well as prophylactic treatments against IR-induced neurocognitive injury. *In vitro* tissue culture models replicate some aspects of cell and tissue-level IR-induced CNS injury including blood brain barrier disruption (Verma, Passerat de la Chapelle et al. 2022) and immune responses (Lojek, Williams et al. 2024) and can be used to investigate the underlying mechanisms.

Space radiation is composed of mixed radiation, a major constituent of which is ionizing charged particles at high energies of up to GeV/amu (Papadopoulos, Kyriakou et al. 2023). High energy charged particle radiation is rare on earth and biophysically distinct from more common terrestrial IR sources such as ionizing photons including X-ray and gamma radiation or common low-energy ionizing particles such as electrons and alpha particles. Importantly, biological injury imparted by densely ionizing particle radiation and sparsely ionizing photon radiation are similar in outcome, as they both cause single and double stranded DNA breaks, but distinct due to the different linear energy transfer properties of each radiation type. For biological experimentation, sources of particle radiation such as simulated galactic cosmic radiation (GCRsim) are often prohibitively difficult to access and therefore more readily available photon sources of X-ray or gamma radiation are often substituted. While the relative biological effectiveness of gamma radiation is less than that of space radiation, it is the closest reasonable approximation that can be used for evaluating chronic injury, as particle radiation sources including cyclotrons and large particle accelerators are generally not able to provide beam time continuously for more than a few hours due constraints on availability (cyclotrons for example are often needed for radiation therapy), instrument duty cycles, and facility costs. Further, many studies that seek to emulate space radiation injury apply acute IR exposures to match cumulative estimated mission-relevant doses; however, deep space exploration will involve persistent low dose-rate exposure over the course of months. While acute IR exposure may often be the only practical and attainable means of simulating space radiation in many in vitro studies, documented differential responses due to dose rate (Osipov, Klokov et al. 2004, Matsuya, McMahon et al. 2018, Nair, Engelbrecht et al. 2019, Ma, Li et al. 2024) could alter experimental outcomes.

For decades, animal models, mainly rodents, have been used to understand the effects of radiation on the CNS and have provided much of the mechanistic and behavioral insights regarding IR and CNS injury. Numerous studies report that IR exposure can alter rodent behavior and lead to neurological deficits associated with adverse neurological conditions in humans. For example, simulated galactic cosmic ray irradiation (GCRsim; lunar-relevant doses of 5 to 50 cGy) induces changes in mouse behaviors associated with hippocampal and medial prefrontal cortex circuitry (Puukila, Siu et al. 2023), disruption of high-cognitive tasks (Stephenson and Britten 2022), and structural alterations in brain tissue at protracted timepoints (>5 month) (Alaghband, Klein et al. 2023). However, such studies which report behavioral changes fail to provide mechanistic insight on cell and tissue level injury. Similarly, space flight studies using rodents on the international space station (ISS) report changes to rodent brain physiology (Latchney, Rivera et al. 2014); however, these studies rely on comparatively short (< 1 month) exposure to the space environment, and necessarily investigate the combined exposure to space radiation, gravitational changes and other spaceflight stressors. In addition, space radiation exposure on the ISS has significantly lower relative biological effectiveness compared to deep space radiation beyond the Earth’s magnetic field. (Jones, Pietrzyk et al. 2008). Thus, the biological effects of radiation dose rate remain to be analyzed in a comprehensive, mechanistic manner with a focus on CNS cell-type specific injury and functional phenotypes.

Traditional radiation biology studies, which use lower model organisms to uncover fundamental molecular IR-induced injury, have shown that dose rate affects DNA damage (Vilenchik and Knudson 2006, Nair, Engelbrecht et al. 2019) and studies using rodent models have observed differential responses to acute and chronic IR exposures (Takahashi, Misumi et al. 2020, Alaghband, Klein et al. 2023). Similarly, differences in non-CNS *in vitro* models show that the functional effects, such as immune response and cytoskeletal organization, are unequal between cumulative equivalent acute and chronic IR doses (Tavakol, Nash et al. 2024). Meanwhile, *in vitro* models of the CNS, likely owing to the difficulty of prolonged experiments, have primarily been studied following acute (< 2 h) rather than chronic (> 2 days) IR exposure. Thus, fundamental cell and tissue-level functional changes in CNS immune response and electrophysiology due to chronic IR exposure remain to be explored. To address this knowledge gap, it is essential to develop a system for chronic irradiation of CNS *in vitro* cultures compatible with longitudinal functional readouts.

Identifying drugs that protect against the CNS effects of space radiation will likely be key to ensuring safe and successful missions to the Lunar or Martian surfaces. The development of tissue culture models which recapitulate essential aspects of IR induced injury would serve as a key platform for evaluating prospective radioprotectant drugs. Much existing research has been done to evaluate the radioprotective properties of various compounds; however, many studies utilize rudimentary tissue culture platforms that do not recapitulate complex aspects of the CNS and/or are anchored on traditional readouts of IR injury (such as DNA damage) that provide minimal relevance to fundamental CNS functions (Smith, Kirkpatrick et al. 2017). Amifostine is a drug which has received limited approval for specific instances of acute radiation syndrome (Singh and Seed 2019). This drug and others like it represent candidate radioprotectants which merit evaluation for neuroprotective properties.

To date, 2D and 3D *in vitro* CNS models have found limited application in assessing space radiation pertinent injuries (Verma, Passerat de la Chapelle et al. 2022, Oyefeso, Goldberg et al. 2023, Roggan, Kronenberg et al. 2023); these few articles generally report mild reactivity in astrocytes as a product of space radiation-related stress. Relevant *in vitro* studies which incorporate CNS cell types include the use of astrocyte monocultures to assess inflammatory phenotypes (Roggan, Kronenberg et al. 2023), organ-on-chip cultures systems which examine vascular permeability and astrocyte reactivity (Verma, Passerat de la Chapelle et al. 2022), and human cerebral organoids to measure transcriptional phenotypes and cell-type specific phenotypes (Oyefeso, Goldberg et al. 2023). Existing space radiation-relevant in vitro studies (1) are limited in number, (2) rarely incorporate functional biological readouts such as assessment of neuroimmune and electrophysiologic phenotypes, which may be more directly related to neurobehavioral impairments, and (3) do not assess injury in chronic exposure scenarios. Looking ahead, CNS *in vitro* models designed for emulating the unique properties of space radiation exposure and examining important functional injury phenotypes will inform critical approaches to space radiation injury mitigation in CNS tissue.

Here, we utilized an engineered 3D collagen hydrogel-based cortical tissue model (CTM) alongside traditional 2D cultures to characterize electrophysiology and neuroimmune phenotypes induced by acute and chronic gamma radiation at Mars mission-relevant doses with and without pre-treatment with the radioprotectant drug amifostine. We examine multiple molecular and cellular outcomes: DNA damage and repair; immune response based on neuronal and glial reactivity; secreted cytokines and chemokines; and network level electrophysiology. We compare IR responses to a standard radiomimetic (bleomycin) based on microelectrode array (MEA) electrophysiology in 2D cultures. Overall, we provide a dose-rate analysis of multiple IR outcomes at molecular, cellular and tissue level, to serve as a benchmark for understanding CNS injury from space radiation.

## Methods

### CTM preparation and co-culture maintenance

Cellularized collagen hydrogels were prepared as previously described (Ghezzi, Marelli et al. 2012, Lojek, Williams et al. 2024). Briefly, acid solubilized type I rat tail collagen (2.05 mg/ml, First Link Ltd., UK) was mixed 4:1 with acid neutralizing cell suspension solution and 80:1 with stem cell qualified ECM solution (CC131-5ML, Sigma Aldrich, MA). The solution was transferred to a 24 well plate (1 ml/well) and incubated at 37 °C for 75 min to polymerize. Final cell densities were estimated to be 10^6^ cells/well. Following polymerization, each gel underwent a plastic compressive force of 1 kPa applied for 5 min at room temperature to create high-density collagen constructs and biopsy punched (33-36, Integra, NH) into 6 mm diameter CTMs. Cultures (both 2D and 3D) were maintained using a culture medium of DMEM with Glutamax (10567022, Thermo-Fisher, MA) with 2% B27 (17504044, Thermo-Fisher, MA), 8 µg/mL L-ascorbic acid (A8960-5G, MilliporeSigma, MA) and 1% Pen-Strep (15140122, Thermo-Fisher, MA). 10% fetal bovine serum (26140079, Thermo-Fisher, MA) was included in the culture medium on the day of seeding but omitted for every feeding thereafter. Medium exchanges were carried out by removing 50% of the media volume and replacing with an equivalent amount of fresh medium three times per week until the beginning of experimental treatment (Figure 1).

**Figure 1.**
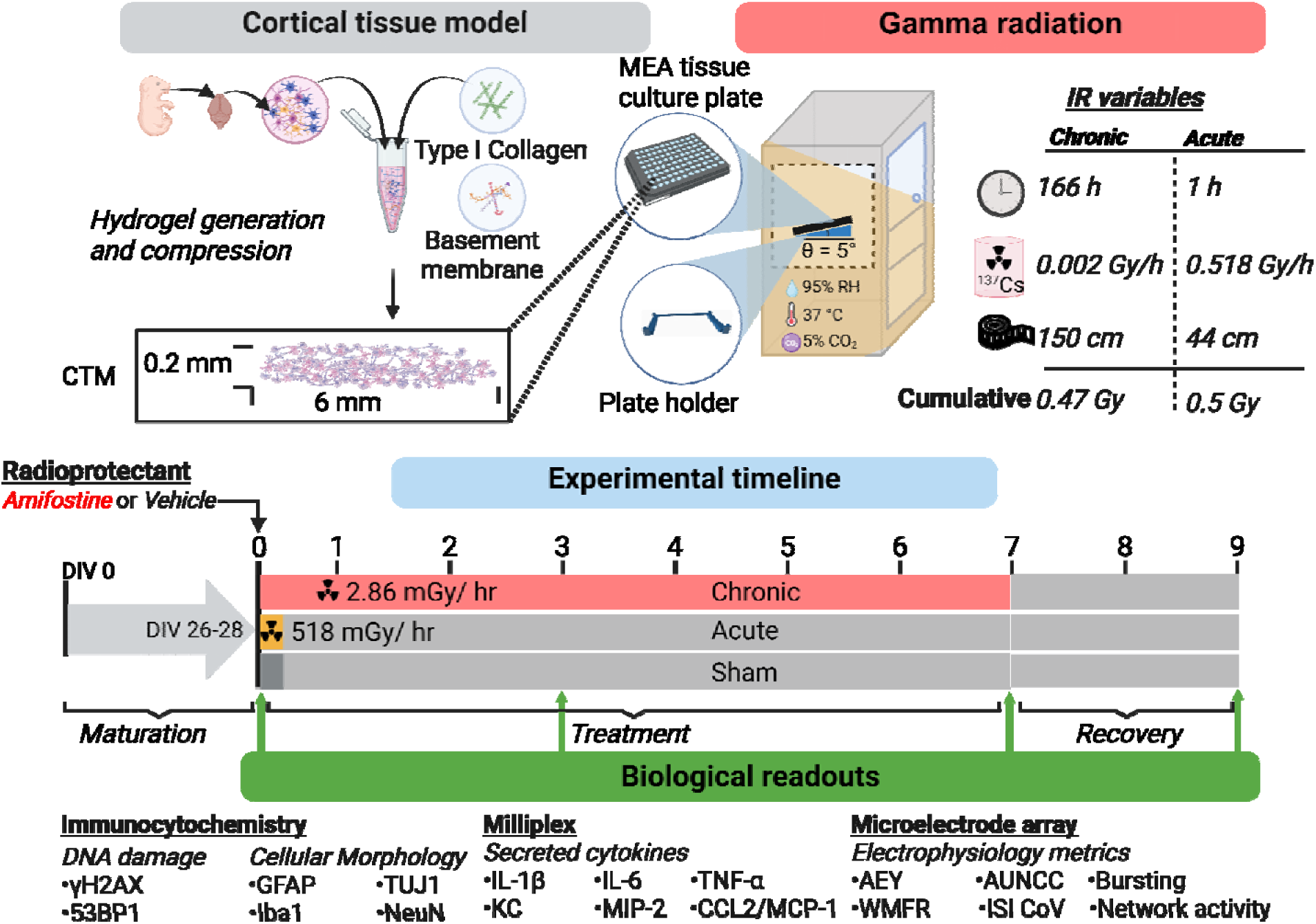
Experimental workflow of CTM gamma radiation exposure. CTMs were prepared by seeding mECNs into type I collagen solution enriched with basement membrane proteins. CTMs were manufactured by an established plastic compression process which produced a dense tissue-like cell-seeded construct. A custom portable tissue culture incubator allowed for CTMs and 2D cultures to be maintained under controlled temperature (37°C), Humidity (>95% RH), and CO_2_ content (5%) while undergoing gamma radiation exposure for up to 7 days. Acute exposure was achieved by placing cultures in the tissue culture incubator at 44cm from a high activity Cs source for 1 h to achieve a dose rate of 0.518 Gy/h and chronic irradiation was achieved by placing cultures at 150 cm from a low activity Cs source to achieve a dose rate of 0.00286 Gy/h for ∼7 days. Sham treated samples experience the same conditions without gamma radiation exposure. Cultures were matured to DIV 26-28 prior to experimentation. Culture media samples were retained from a media change immediately prior to experimentation, and at 3, 7, and 9 days following the start of exposure. Chronically irradiated samples underwent 2 days of recovery under normal tissue culture conditions after the end of gamma radiation exposure. Biological readouts include DNA damage, cytomorphology, and immune reactivity by ICC, cytokine secretion by milliplex assay, and network level electrophysiology metrics recorded from 2D cultures by MEA.

### 2D MEA culture and maintenance

In the case of 2D cultures, 48-well MEA plates (Axion Biosystems, GA) were pretreated with 0.1% Poly(ethyleneimine) PEI and 20 µg/ml laminin (20 ng/ml, Sigma Aldrich, MA). 5 µl of dissociated murine embryonic cortical neuronal tissue (mECN) cell solution was placed at the center of each well and allowed to incubate for at least 20 min. After visualizing cell attachment under a benchtop phase microscope, wells were flooded with 200 µl of complete growth medium.

### Embryonic mouse cortical dissection

Embryonic mouse cortical dissection and 2D culture was carried out as previously described (Black, Mondal et al. 2013). All animal procedures were conducted according to UML IACUC approved protocols (Protocol Number: 22-10). A full-time veterinarian participates in the animal care program at UML. Veterinary care includes a program for prevention of disease, daily observation, and surveillance for assessment of animal health, appropriate methods of disease control, diagnosis and treatment, guidance of animal users in appropriate methods of handling and restraint, anesthesia, analgesia, and euthanasia, and monitoring of surgical programs and post-surgical care, all in compliance with AVMA guidelines. The study is reported in accordance with ARRIVE guidelines. Briefly, timed-pregnant female mice (E18 – 022TP-Charles River Laboratories, Inc - Wilmington, MA, USA) were euthanized by cervical dislocation and cesarean section was performed to expose and isolate embryonic sacs. Each embryo was removed, decapitated, and its head washed twice in ice-cold HBSS. Cortical lobes were isolated, minced with surgical scalpel blade, and pooled in dissociation solution (0.025% Trypsin and 2500 U/ml DNase (11284932001, Millipore Sigma, MA)) for approximately 25 min with intermittent mechanical trituration using a 1 ml pipet until homogenized (approximately 30 times). Dissociation was quenched with medium containing 10% FBS and the cell solution was centrifuged at 300xg for 5 min. The resulting cell pellet was resuspended in the appropriate volume, depending on whether the application was 2D or 3D cultures, and seeded onto MEA plates or in 3D constructs. All protocols involving animals were carried out in accordance with institutional guidelines in compliance with the Guide for the Care and Use of Laboratory Animals (ISBN-13: 978-0-309-15400-0, National Academy Press 2011). UMass Lowell Campus maintains the Public Health Service Policy Assurance number A3867-06.

### CTM gamma radiation exposure and radioprotectant drug treatment

Amifostine is a radioprotectant drug which is approved by the FDA for treatment of acute radiation syndrome in specific radiation exposure scenarios (Singh and Seed 2019) (King, Joseph et al. 2020). CTM and 2D mECN cultures were pre-treated once with 250 µM amifostine or vehicle (water) 1 h prior to gamma radiation or sham exposure in a 50% media exchange. Mature cultures (DIV 26-28) were treated with either acute or chronic gamma radiation from ^137^Cs sources at the UMass Lowell Radiation Laboratory. A portable tissue culture incubator compatible with the spatial constraints of the source room was built by modifying a small temperature-controlled incubator (Mytemp Mini H-2200, Marshall Scientific, NH) to accommodate continuous flow of a pre-mixed 95% air 5% carbon dioxide (X02AI95C2000117, Airgas, PA). For all acute doses tissue culture plates were positioned in the incubator 44 cm from a high activity ^137^Cs NIST traceable source for a dose rate of 0.518 Gy/h with a maximum source variability of 6.46%. Samples were exposed for 58 minutes to achieve a total cumulative dose of 0.5 Gy. For chronic treatment tissue culture plates were positioned in the incubator at 150 cm from a low activity ^137^Cs NIST traceable source for a dose rate of 0.0028624 Gy/h with a maximum source variability 6.89%. Cultures treated chronically were irradiated in two identical experiments occurring at different times for cumulative total times of 166.61 and 166.88 h giving a total cumulative dose of 0.47690 Gy and 0.47768 Gy, respectively. For chronic cultures the radiation source was shut down briefly for 4 h on the 3^rd^ day of the treatment to facilitate media exchanges and MEA recordings. All cultures (sham, acute, and chronic) were maintained for an additional 2 days following the end of chronic irradiation (at day 7), for a total of 9 days from the beginning of acute, chronic, or sham exposure (Figure 1). To minimize self-attenuation from the multi-well plate itself, all plates, including the sham-treated plates, were elevated at a 5° angle with respect to the incubator shelf for the duration of the chronic irradiation period (7 days) using a custom-built 3D printed wedge. Both acute and chronic gamma radiation configurations were in an open beam geometry. Positional differences between the wells closest to the ^137^Cs source and those furthest away constitute a 16.587 % error for the acute exposure (high activity source) and a 4.478 % error for the chronic exposure (low activity source). Gamma radiation dose rates inside of the tissue culture incubator were independently verified for each source using an ion chamber survey meter (RO-20, Thermo Scientific, MA).

### Immunocytochemistry, imaging, and analysis

Immunocytochemistry (ICC) preparation, imaging, and analysis for DNA damage markers, glial reactivity, and cytomorphology were carried out as previously described (Lojek, Williams et al. 2024). Briefly, CTM samples were fixed by incubating in room-temperature 4% paraformaldehyde (AAJ61899AK, Thermo-Fisher, MA) for 60 min, followed by 3x washes with 1x phosphate buffer saline (PBS; 70011069, Thermo-Fisher, MA) and stored at 4 °C until antibody labeling and imaging. Samples were permeabilized and blocked by incubating 30 min in 0.2% Triton X-100 in PBS (Sigma-Aldrich, MO) followed by 60 min in 1% bovine serum albumin (Sigma-Aldrich, MO) in PBS. Samples were washed 3x in PBS and then incubated with primary antibodies overnight, washed 3x with PBS and incubated secondary antibodies for at least 4 h, followed by another 3x wash with PBS, 20 min incubation with DAPI, and the final 3x wash with PBS less than 1h prior to imaging.

The following antibodies were used for ICC analyses: <u>53BP1</u>: mouse anti-53BP1 (1:1000) (MA538649, Thermo Fisher, MA) and goat anti-mouse IgG1 secondary antibody (1:1000 concentration) (A-21126, Thermo Fisher, MA). <u>Neurites</u> (βIII-tubulin): Chicken anti-βIII-tubulin (Tuj1) (1:1000) (NB100-1612, Novis Biological) and goat anti-chicken (1:1000) (A32933, Thermo Fisher, MA). <u>Neuronal nuclei</u> (NeuN): Rabbit anti-NeuN (1:1000) (ab177487) and Donkey anti-rabbit (1:1000) (A-21206 Thermo Fisher, MA). <u>Astrocytes</u> (GFAP): Chicken anti-GFAP (1:1000) (PA1-10004, Thermo Fisher, MA) and goat anti-chicken (1:5000) (A32933, Thermo Fisher, MA). <u>Microglia</u> (Iba1): Rabbit anti-Iba1 (1:1000) (AB178846, Abcam, MA) and donkey anti-rabbit IgG antibody (1:1000) (A21206, Thermo Fisher, MA).

For each sample, at least three regions of interest were captured at either 20 or 40x magnification as z-stacks using a Leica SP8 confocal laser scanning microscope (CLSM, Leica Biosystems, MA) in line-sequence mode to reduce fluorescence emission overlap. Image masking and quantification of mean intensities was carried out as previously detailed (Lojek, Williams et al. 2024). Quantification of neurite length, number and size/circularity of glia, glial activation marker intensity, number of NeuN-associated nuclei, and DNA damage marker intensity was carried out using custom boutique macros in ImageJ (NIH, USA).

### Multiplexed cytokine assay

A multiplex cytokine assay kit (MCYTOMAG-70K, MilliporeSigma, MA) targeting pro-inflammatory cytokines Interleukin-1 beta (IL-1β), Interleukin 6 (IL-6), KC/CXCL1, Monocyte Chemoattractant Protein 1 (MCP-1/CCL2), Macrophage Inflammatory Protein 2 (MIP-2), and Tumour Necrosis Factor alpha (TNF-α) was used in conjunction with a Luminex SD liquid handling device and Luminex 200 plate reader in accordance with the product’s protocol. Medium samples were collected from CTMs 7 days following IR exposure, centrifuged at 300xg to remove any residual cells/debris, and stored at -80 °C (< 4 months). Analyte concentrations were calculated by comparing median fluorescence intensities with associated standard curves using Belysa® Immunoassay curve fitting software (Merck Millipore, MA) as previously described (Adelfio, Martin-Moldes et al. 2023). Resulting protein concentrations obtained from multiplex analysis were normalized by their respective estimated cell number from CLSM analysis of DAPI nuclei-stained samples.

### Extracellular electrophysiology recordings

Substrate integrated MEA recordings were carried out as previously described (Black, Atmaramani et al. 2017, Black, Atmaramani et al. 2018, Black, El Ghazal et al. 2024). Briefly, extracellular action potentials (EAPs) were identified by applying an adaptive ±5.5σ threshold to filtered continuous recordings (12.5 kHz sampling rate) performed using the Axion Maestro Pro (Axion Biosystems, GA). Baseline and post-treatment recordings consisted of a 30 min acclimation period followed by a 10 min recording, while maintaining samples at 37 °C and 5% CO_2_. Weighted mean firing rates (WMFR), active electrode yields (AEY), area under the normalized cross correlogram (AUNCC), interspike interval coefficient of variance (ISI CoV), and intrinsic and network burst frequency (NBF) were quantified using Axion’s Neural Metrics Tools. Statistical comparisons were carried out based on difference over sum (DoS) normalized values as compared to baselines.

### Statistical Analysis

R Studio packages tidyverse, ggpubr, and rstatix were used for data processing, plotting, and statistical analysis, respectively. Subsequent to either Shapiro–Wilk or Kolmogorov–Smirnov normality test, a Student’s T test for equal variances/normal distribution or Mann-Whitney U-test for unequal variances was used to calculate p-values for the associated groups when comparisons between two groups were made, as in the case of DNA damage assessments. For comparisons across more than two conditions, as for cellular reactivity, cytomorphology, cytokine secretion, and MEA, a 2-way-ANOVA test with a subsequent Holm-Bonferroni post-hoc test was used. To maintain clarity, when comparing across multiple groups, post-hoc statistical differences were retained for pairs that reflect the effects of radiation or drug alone Acute-Amifostine & Acute-Vehicle, Chronic-Amifostine & Chronic-Vehicle, Sham-Amifostine & Sham-Vehicle, Acute-Vehicle & Sham-Vehicle, Chronic-Vehicle & Sham-Vehicle, Chronic-Vehicle & Acute-Vehicle, Acute-Amifostine & Sham-Amifostine, Chronic-Amifostine & Sham-Amifostine, Chronic-Amifostine & Acute-Amifostine. Complete tables containing all statistical comparisons are included in the supplementary information. Values of p<0.05 were considered statistically significant.

## Results

### DNA damage assessment by 53BP1

IR is well understood to cause cytotoxicity by inducing various types of DNA damage, including single strand breaks (SSB), double strand breaks (DSB), formation of abasic sites and cross-links. This type of injury is highly deleterious and can result in genetic mutations, gene deletion, and/or genetic translocation (Garte and Burns 1991, Iliakis, Wang et al. 2003, Mavragani, Nikitaki et al. 2019). The biochemical pathways involved in DNA DSBs and their repair are well characterized (Penninckx, Pariset et al. 2021). After exposure to acute or chronic gamma radiation, we assessed DNA damage in cortical models by quantifying the DNA repair protein 53BP1 using immunocytochemistry (ICC) staining and quantitative CLSM. Representative 53BP1 immunostained images of vehicle-treated samples at 30 min following acute gamma radiation exposure at 0.5 Gy alongside time-matched sham controls are shown in Figure 2A as well as chronic treated samples after 3 and 7 days (cumulative doses 0.2 and 0.47 Gy respectively). Supplemental Figure 1 shows representative images of amifostine-treated samples stained for 53BP1.

**Figure 2.**
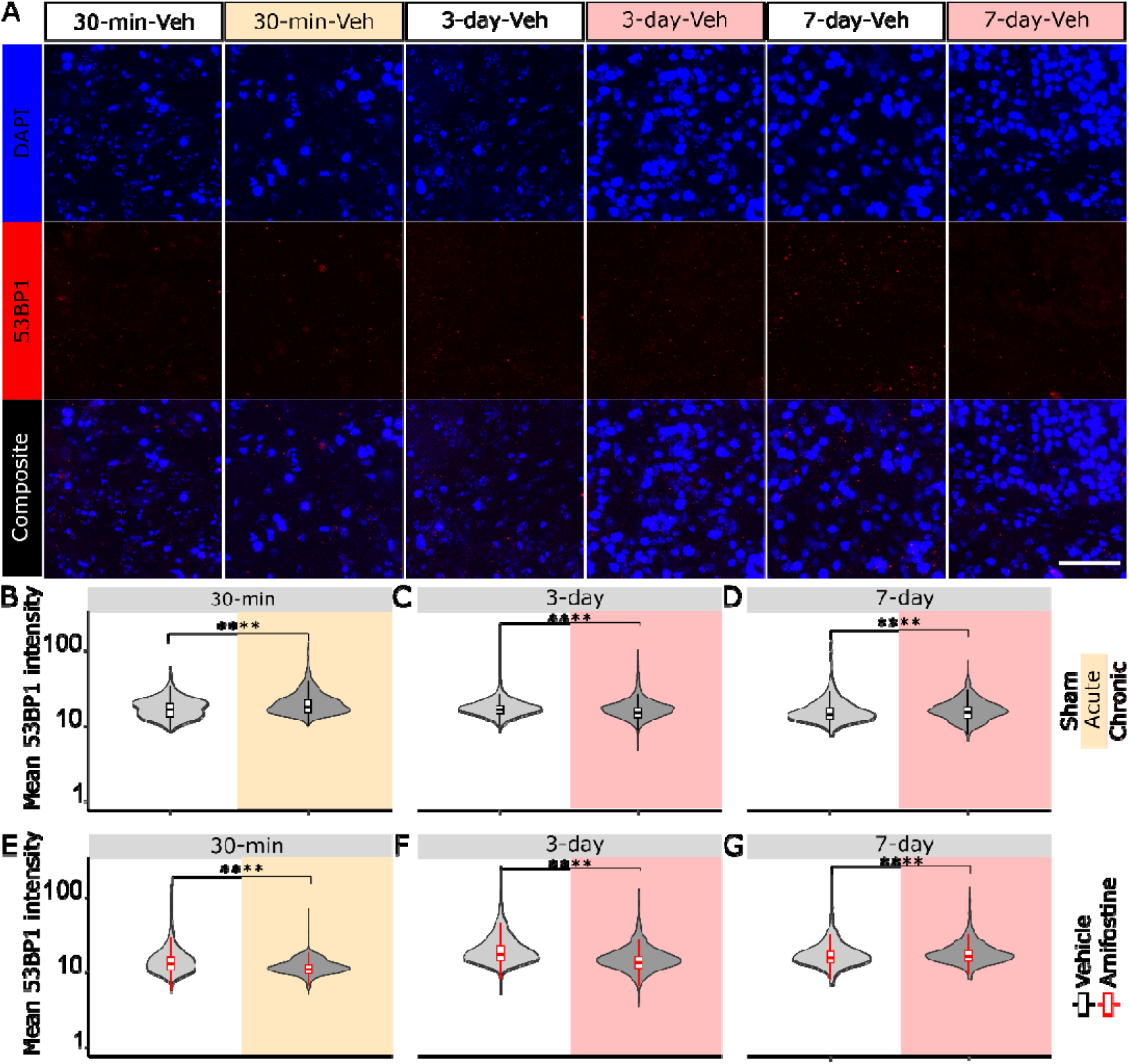
DNA damage quantification within CTM due to gamma radiation. A. Representative CLSM images of DAPI (blue) and 53BP1 (red) ICC staining in sham (white) acute (light yellow) and chronic (light red) cultures. Scale bar = 50 µm. B. Quantification of DNA damage marker 53BP1 mean intensity per nuclei object at 30 min post-acute exposure and C. 3-days and D. 7-days after the beginning of chronic exposure in vehicle treated samples (black outline) and at E. 30-min post-acute exposure, F. 3-days and G. 7-days following the beginning of chronic exposure with amifostine treatment (red outline).

Quantitative analysis of 53BP1 marker intensity shows a significant increase in acute-vehicle (20.039 ± 9.346) relative to sham-vehicle samples (16.912 ± 4.902) at 30-min after 0.5 Gy acute exposure (p<0.0001) (Figure 2B), a significant decrease in chronic-vehicle (15.611 ± 5.507) relative to sham-vehicle (16.707 ± 5.407) at 3-days after the beginning of exposure (0.2 Gy cumulative dose) (p<0.0001) (Figure 2C), and a significant increase in chronic-vehicle (15.717 ± 4.990) relative to sham-vehicle (15.405 ± 5.574) at 7-days after the beginning of exposure (0.47 Gy cumulative dose) (p<0.0001) (Figure 2D). Analysis of radioprotectant and vehicle treated samples shows a significant decrease in acute-amifostine (11.219 ± 2.634) relative to sham-amifostine samples (14.552 ± 7.874) at 30-min after 0.5 Gy acute exposure (p<0.0001) (Figure 2E), a significant decrease in chronic-amifostine (14.497 ± 6.066) relative to sham-amifostine (20.046 ± 10.539) at 3-days after the beginning of exposure (0.200 Gy cumulative) (p<0.0001) (Figure 2F), and a significant increase in chronic-amifostine (17.734 ± 7.018) relative to sham-amifostine (17.453 ± 8.457) at 7-days after the beginning of exposure (p<0.0001) (Figure 2G).

### Assessment of cytomorphology and immune reactivity

An assessment of cytomorphology was used to evaluate functional changes to cell-seeded CTMs treated with acute, chronic, or sham radiation following pre-treatment with amifostine or a vehicle. Representative maximum projection images of ICC staining show DAPI (blue), the astrocyte marker GFAP (red) and the neuronal marker NeuN (green) fixed at 9-days after the beginning of exposure (Figure 3A).

**Figure 3.**
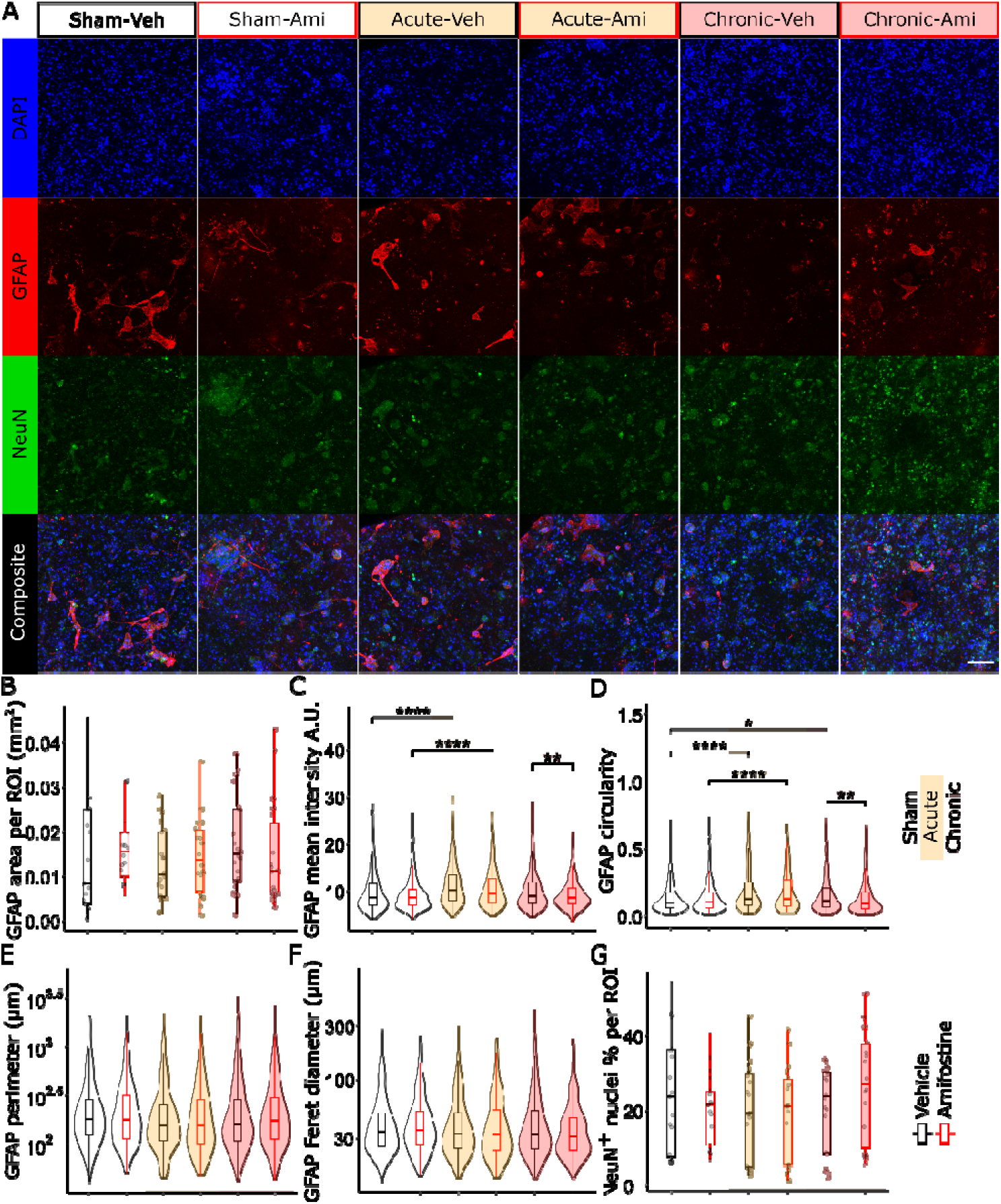
Astrocyte reactivity, cytomorphological, and neuronal nuclei population within CTMs following acute and chronic gamma radiation. A. representative CLSM images of DAPI (blue) GFAP (red), and NeuN (green) staining in CTMs shown as maximum intensity projections from ICC at 9 days following the beginning of acute (yellow shade), chronic (red shade), or sham (no shading) gamma radiation exposure with amifostine (red outline) or vehicle (black outline) pre-treatment. (Scale bar = 100 µm). Quantification of mean GFAP intensity per cell object (B), GFAP total area (C) GFAP cell object mean intensity (D) circularity (E), perimeter (F) ferret diameter (G), and percent of nuclei objects positive for the neuronal marker NeuN per ROI.

Total GFAP area per ROI, a measure of glial reactivity, for sham-vehicle (0.014 ± 0.013 mm^2^,n= 19) , sham-amifostine (0.0166 ± 0.008 mm^2^, n= 18), acute-vehicle (0.013± 0.008 mm^2^, n= 20), acute-amifostine (0.0144 ± 0.009 mm^2^, n=18), chronic-vehicle (0.017 ± 0.011 mm^2^ ,n=20), and chronic-amifostine (0.014 ± 0.009 mm^2^, n=18) were not significantly different (Figure 3B). Quantification of mean GFAP intensity per cell object showed a significant increase in GFAP intensity in the acute-vehicle group (11.206±4.172, n= 412) relative to the sham-vehicle group (9.659± 3.924, n =390) (p< 0.0001), acute-amifostine (10.602 ±4.126, n=433) was increased relative to sham-amifostine (9.179± 3.239, n=535) (p< 0.0001), and chronic-amifostine (9.280 ±3.108, n= 533) was decreased relative to chronic-vehicle (10.009± 3.809, n= 533) (p= 0.0096) (Figure 3C).

GFAP cell object morphology was assessed by measuring circularity, perimeter, and Feret diameter. Cell circularity, which when increased indicates inflammation in immune cells (Hu, Huang et al. 2021), was significantly increased in the acute-vehicle group (0.187± 0.155) and the chronic-vehicle (0.163 ± 0.142) group relative to the sham-vehicle group (0.136± 0.117, n =390) (p< 0.0001 and p = 0.0172 respectively), acute-amifostine (0.179 ± 0.141) was increased relative to sham-amifostine (0.136 ± 0.110) (p< 0.0001), and chronic-amifostine (0.136 ±0.125) was decreased relative to chronic-vehicle (0.163± 0.142) (p= 0.0075) (Figure 3D). Cell object perimeter for sham-vehicle (280.367± 305.789 µm), sham-amifostine (269.095± 274.267 µm), acute-vehicle (236.542± 247.681 µm), acute-amifostine (238.818± 239.341 µm), chronic-vehicle (266.153± 310.675 µm), and chronic-amifostine (267.987± 297.104 µm) were not significantly different (Figure 3E). Cell object Feret diameter for sham-vehicle (46.432 ± 38.180 µm), sham-amifostine (45.336 ± 31.705 µm), acute-vehicle (43.925 ± 33.697 µm), acute-amifostine (43.968 ± 34.007 µm), chronic-vehicle (46.403 ± 42.725 µm), and chronic-amifostine (45.336 ± 31.705 µm) were not significantly different (Figure 3F).

To assess the number of neuronal cells in CTMs at 9-days following the beginning of exposure the number of NeuN^+^ DAPI nuclei were measured. NeuN+ nuclei for sham-vehicle (24.5 ± 16.1 %, n= 15598 nuclei), sham-amifostine (20.4 ± 9.5 %, n= 18509 nuclei), acute-vehicle (20.3 ± 13.3 %, n= 22364 nuclei), acute-amifostine (19.2 ± 12.7 %, n= 24859 nuclei), chronic-vehicle (20.9 ± 11.4 %, n= 24033 nuclei), and chronic-amifostine (26.9 ± 15.6 %, n= 23931 nuclei) were not significantly different (radiation p= 0.419, radioprotectant p= 0.904, radiation: radioprotectant p= 0.240) (Figure 3F) . Additionally, mean DAPI nuclei per mm^3^ for sham-vehicle (15546.9 ± 9447.5, n= 15598 nuclei), sham-amifostine (19473.3 ± 5653.6, ROI = 19), acute-vehicle (21176.2 ±13241.9 , ROI= 20), acute-amifostine (26154.1 ± 15239.5, ROI= 18), chronic-vehicle (22756.5 ± 11406.3, ROI= 20), and chronic-amifostine (25177.7 ± 13957.3, ROI= 18) were found no significant between relevant groups by post-hoc tests (Supplemental Figure 2). A complete table of all significant pairwise comparison values can be found in supplemental information table 1.

Representative maximum projection images of ICC staining show DAPI (blue) the neuronal marker βIII-tubulin (red) and the microglia marker Iba1 (green) at 9-days following the beginning of exposure (Figure 4A). Iba1 total area per ROI, a measure of glial reactivity, for sham-vehicle (0.0085± 0.01578 mm^2^, n=9) , sham-amifostine (0.0022 ± 0.0035 mm^2^, n=7), acute-vehicle (0.0015± 0.0021 mm^2^, n=11), acute-amifostine (0.0023± 0.0051 mm^2^, n=12), chronic-vehicle (0.0031± 0.0037 mm^2^ ,n= 9), and chronic-amifostine (0.0022± 0.0015 mm^2^, n=7) were not significantly different across groups (Figure 4B). Quantification of mean Iba1 intensity per cell object showed a significant reduction in sham-amifostine (7.226 ± 1.627, n= 154) samples relative to sham-vehicle samples (8.044 ± 1.905, n= 320) (p< 0.0001), additionally both acute-vehicle (7.311 ± 1.598, n= 153) and chronic-vehicle (2.051 ± 126.270, n=218) samples were significantly decreased relative to sham-vehicle (p= 0.0003 and p= 0.0071 respectively), acute-amifostine (6.963 ± 1.126, n= 143) and chronic amifostine (1.700 ± 80.088, n= 196) showed no significant difference compared to sham-amifostine (Figure 4C). The number of Iba1 cell objects per ROI was not significantly different across groups (Supplemental Figure 3).

**Figure 4:**
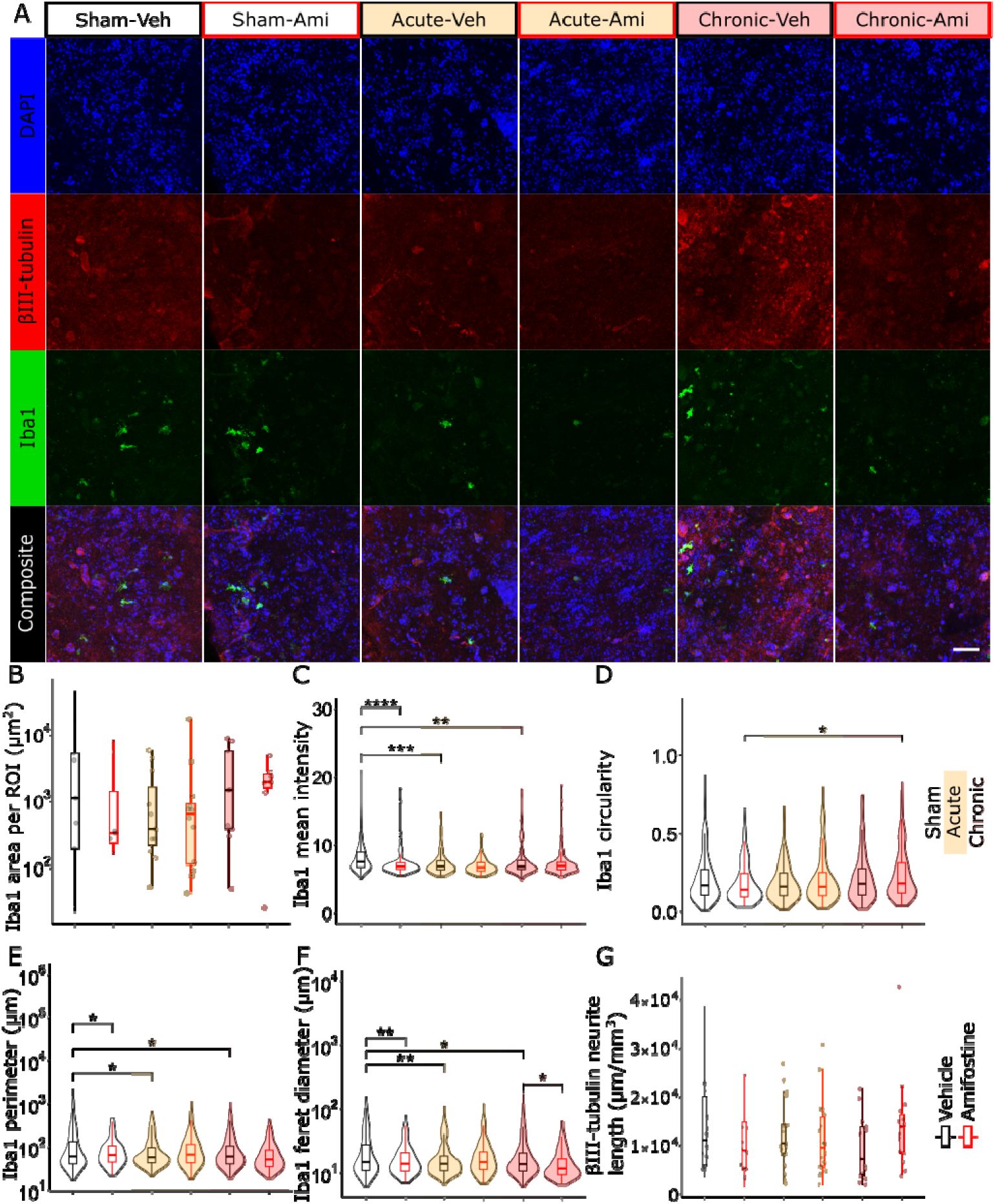
Microglia reactivity, cytomorphology, and neurite morphology in CTMs following to acute and chronic gamma radiation. A. representative CLSM images of DAPI (blue) Iba1 (green), and βIII-tubulin (red) staining in CTMs shown as maximum intensity projections from ICC at 9 days following the beginning of acute (yellow shade), chronic (red shade), or sham (no shading) gamma radiation exposure with amifostine (red outline) or vehicle (black outline) pre-treatment. (Scale bar = 100 µm). B. Quantification of mean Iba1 intensity per cell object, C. circularity, D. Feret diameter, E. perimeter, and F. summed βIII-tubulin neurite length per ROI.

Iba1 cell object circularity was measured for sham-vehicle (0.210 ± 0.149), sham-amifostine (0.188 ± 0.137), acute-vehicle (0.187 ± 0.120), acute-amifostine (0.208 ± 0.157), chronic-vehicle (0.218 ± 0.156), and chronic-amifostine (0.237 ± 0.158). Significant pairwise comparisons were found between sham-amifostine and chronic-amifostine samples (p= 0.0314) (Figure 4D). Iba1 cell object perimeter was found to be increased in acute-vehicle (90.916 ± 94.380 µm), chronic-vehicle (97.841 ± 118.470 µm), and sham-amifostine (90.411 ± 82.619 µm) groups relative to sham-vehicle (137.800 ± 231.080 µm) group (p= 0.0159, p= 0.0245, and p= 0.0148 respectively). Acute-amifostine (103.300 ± 128.156 µm) and chronic-amifostine (70.833 ± 60.240 µm) were not found to be significantly different from other relevant groups by pairwise comparison (Figure 4E). Iba1 cell object Feret diameter was found to be significantly decreased in acute-vehicle (18.509 ± 14.527 µm), chronic-vehicle (19.696 ± 18.618 µm), and sham-amifostine (17.693± 11.829 µm) groups relative to the sham-vehicle (24.557 ± 24.341 µm) group (p= 0.0063, p= 0.0203, and p= 0.0010 respectively). Chronic-vehicle Feret diameter was significantly increased compared to chronic-amifostine (14.731± 8.998 µm) (p= 0.0422). Acute-amifostine (19.430± 14.790) was not significantly different from any other relevant groups by pairwise comparison (Figure 4F). In summary, cell perimeter and ferret diameter were reduced by acute and chronic radiation, while amifostine increased cell perimeter and decreased ferret diameter. A complete table of all statistically significant pairwise comparison values can be found in supplemental information table 2.

βIII-tubulin, a marker of neuronal processes, was quantified at 9-days following the start of exposure by summing the neurite length in each ROI. For sham-vehicle (13741.457 ± 9847.593 µm/mm^3^), sham-amifostine (10730.406 ± 7411.663 µm/mm^3^), acute-vehicle (12297.082 ± 7214.101 µm/mm^3^), acute-amifostine (11318.334 ± 7850.608 µm/mm^3^), chronic-vehicle (9518.221 ± 6509.855 µm/mm^3^), and chronic-amifostine (14322.838 ± 9414.677 µm/mm^3^) no significantly pairwise differences were found between groups (Figure 4G).

### Endogenous markers secretion

Culture media was collected from both CTMs and 2D cultures prior to acute, chronic, or sham IR exposure and at 7-days following the start of IR exposure. To characterize radiation induced stress, the change in secreted cytokine markers was measured by milliplex assay including IL-1β, IL-6, KC, MCP-1, MIP-2, and TNF-α. Cytokine content in media at 7-days after the start of irradiation was normalized to pre-treatment cytokine measurements for all samples. IL-1β and TNF-α were below the detection limit for both 3D CTM and 2D samples. In 2D cultures, there was no significant change in cytokine content for IL-6, KC, MCP-1, or MIP-2 for any treatment group (n=6) (Figure 5A). In 3D CTM cultures, IL-6 is significantly decreased in Chronic-Vehicle relative to Sham-Vehicle (p= 0.0063), MCP1 is significantly decreased in Amifostine-Vehicle relative to Sham-Vehicle (p= 0.0146), KC is significantly decreased in Chronic-Vehicle relative to Sham-Vehicle (p= 0.0050), and in MIP2 is significantly decreased in Chronic-Vehicle relative to Sham-Vehicle (p= 0.0221)(Figure 5B).

**Figure 5:**
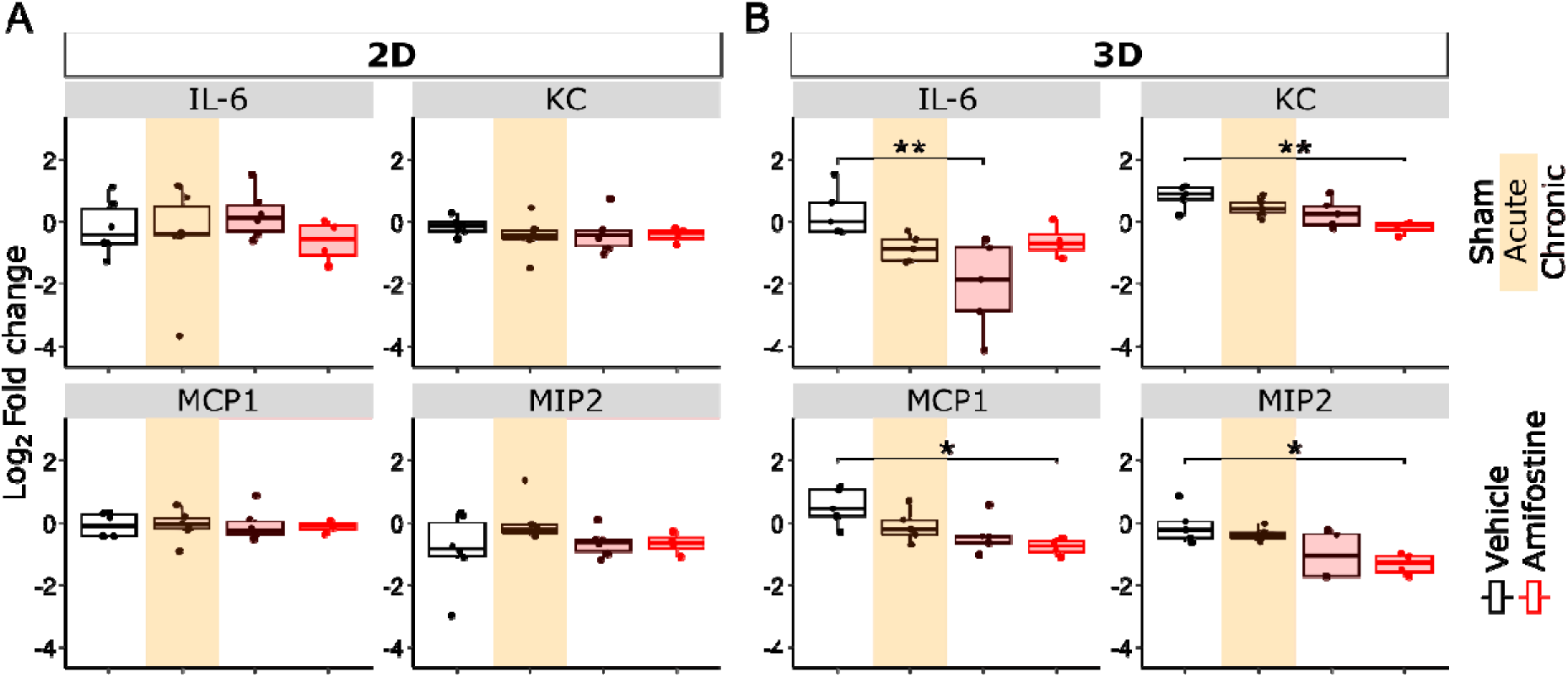
Detection of pro-inflammatory cytokines in tissue culture medium of 2D mECN and 3D CTM cultures. A.) Fold change in secreted pro-inflammatory cytokines IL-6, KC, MCP1, and MIP2 in 2D mECN cultures shown between pre-treatment media and media collected at 7-days following the beginning of gamma radiation exposure, amifostine or both. B.) Fold change in secreted pro-inflammatory cytokines IL-6, KC, MCP1, and MIP2 in 3D CTM cultures shown between pre-treatment media and media collected at 7-days following the beginning of exposure with GR, amifostine or both.

### Electrophysiology after bleomycin treatment

Functional electrophysiology changes in 2D cultures seeded on MEA plates were characterized by comparing relative change between samples treated with the radiomimetic drug bleomycin and those treated with a vehicle. Figure 6A shows phase contrast images of mECN cultures seeded at a high density (50k cells per well) over microelectrodes. A snippet of a representative continuous trace recording is shown (Figure 6B) as well as 60 second representative raster plot which shows the relative time of extracellular action potentials (EAP) for 16 electrodes from a single selected well (Figure 6C). An mECN culture exhibited an average AEY of 82.55% (Figure 6D) and an average WMFR of 10.16 ± 3.87 at DIV 35 (Figure 6E) at which point bleomycin treatment experiments began. All post-treatment MEA recording timepoints (1, 24, and 48 h) for all electrophysiology metrics were normalized by difference-over-sum-normalization to a pre-treatment baseline recording made less than two hours prior to bleomycin treatment. Metrics of electrophysiology including AEY (Figure 6F), WMFR (Figure 6G), AUNCC (Figure 6H), ISI CoV (Figure 6I), number of bursts (Figure 6J), and network burst frequency (Figure 6K) were measured. Plots show the vehicle treatment (black) and a dose curve of bleomycin 0.01 µg/mL, 0.1 µg/mL, 1 µg/mL, 10 µg/mL, and 50 µg/mL (blue gradient). There is a statistically significant reduction of AEY at 48 h (p= 0.0008), WMFR at 48 h (p= 0.0017), AUNCC at 48 h (p= 0.0024), number of bursts at 48 h (p= 0.0027), and NBF at 48 h (p= 0.0346) due to bleomycin at only 50 µg/mL. Results table can be found in supplementary table 3.

**Figure 6:**
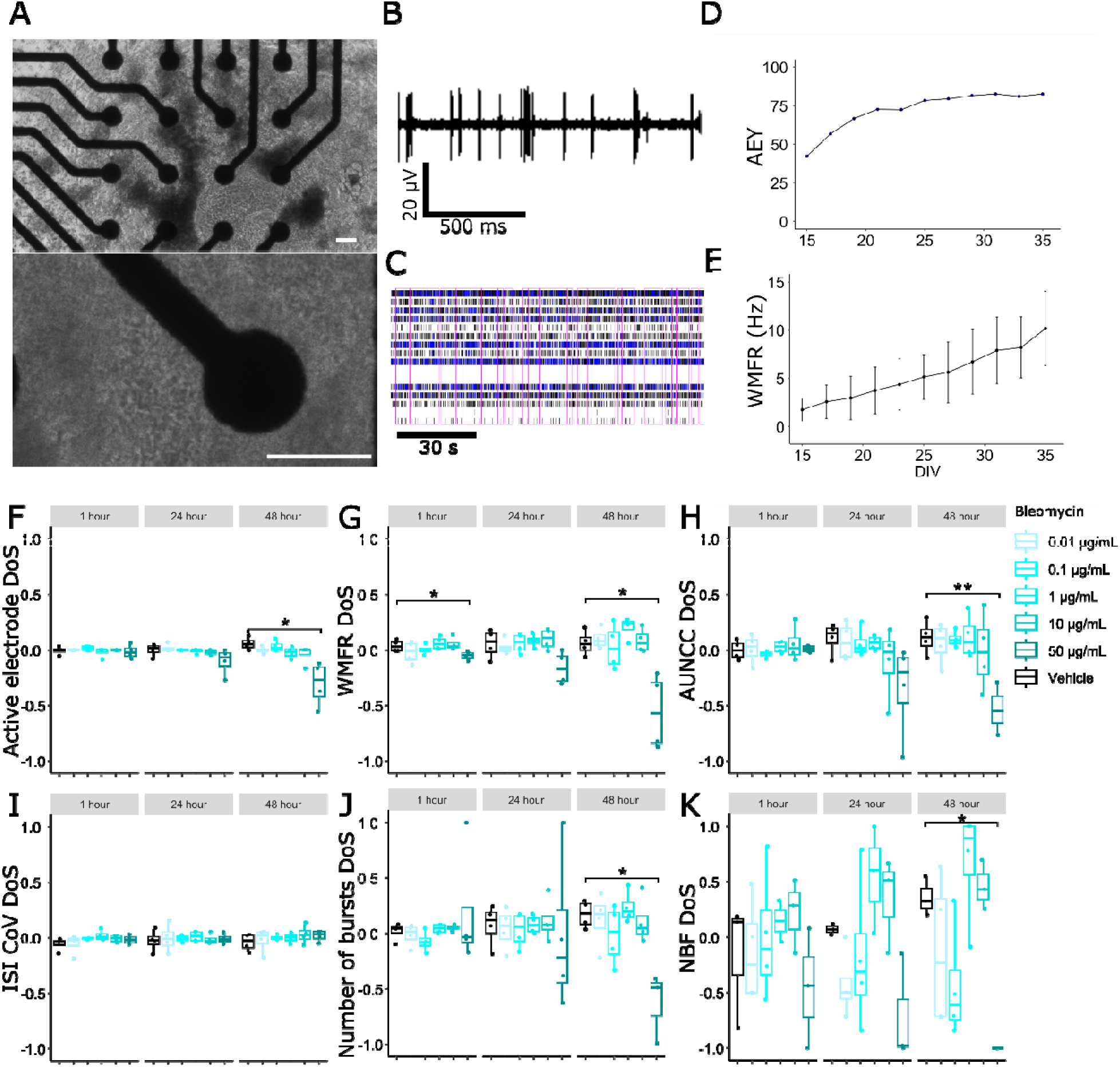
Radiomimetic drug bleomycin effects on electrophysiology. A. Representative phase contrast images of 2D MEA cultures seeded on MEA plates at DIV 25. Scale bar = 50 µm. B. Representative filtered voltage by time for MEA recordings from a single electrode DIV 36. C. Representative raster plot indicating EAP timestamps across all 16 electrodes in the well. Black and blue lines represent single EAPs and intrinsic burst respectively. Pink rectangles highlight network bursts. D. Time progression of 2D culture AE and E. WMFR. Bleomycin effects shown by change relative to pre-treatment timepoint by DoS normalization for AE, F. WMFR, G. AUNCC, H. ISI CoV, I. Number of bursts, and J. NBF. Bleomycin doses shown in blue with vehicle indicated by black.

### Electrophysiology response after sham, acute, and chronic gamma radiation

2D mECN cultures were recorded by MEA to measure electrophysiology phenotypes due to acute and chronic radiation treatment. Three separate mECN cultures exhibited an average AEY of 77.97% and an average WMFR of 10.282 with a standard deviation of between 5.392 and 4.550 at DIV 26-28 at which point IR exposure began. All post-treatment MEA recording timepoints (1 h, days 1 to 9) for all electrophysiology metrics were normalized by difference-over-sum-normalization (DoS) to a pre-treatment baseline recording made less than 2 h prior to radiation, sham, or drug treatment.

Measurement of difference over sum normalized active electrodes indicated significant a reduction due to amifostine in the sham radiation group at 2 through 7-days, in the acute radiation group at 1 through 7-days, and in chronic radiation at 3, 7, 8, and 9-days (Figure 7A). For WMFR statistically significant reductions due to amifostine were measured in the sham radiation group at 2 through 7-days, in the acute radiation group at 2-6 days, and in the chronic radiation group at 3, 7, 8, and 9 days (Figure 7B). For AUNCC there is a significant reduction due to amifostine in the acute radiation group at 3 through 5 days and a statistically significant reduction of acute radiation on AUNCC in amifostine treated groups at 3 and 4 days (p< 0.001, *##*) (Figure 7C). No significant differences were found across any of the relevant groups at any timepoint for or ISI CoV (Figure 7,D). There is a statistically significant reduction in number of bursts due to amifostine in the sham radiation group at 2-6 days, in the acute radiation group at 2-6 days, and in the chronic radiation group at 2 and 7 days (Figure 7E). At day 5, there is a significant reduction in NBF due to acute radiation in the vehicle treated groups, a statistically significant reduction of NBF due to amifostine in the sham radiation group at 2 through 6-days, in the acute radiation group at 2, 3, and 6-days, and no significant effects at any chronic exposure timepoint (Figure 7F). Plots show sham (no shading), acute (yellow shaded), and chronic radiation (pink shaded) with either vehicle (black outline) or amifostine (red outline) pre-treatment. Complete results can be found in supplementary table 4 and all significant pairwise comparison values can be found in supplementary table 5.

**Figure 7:**
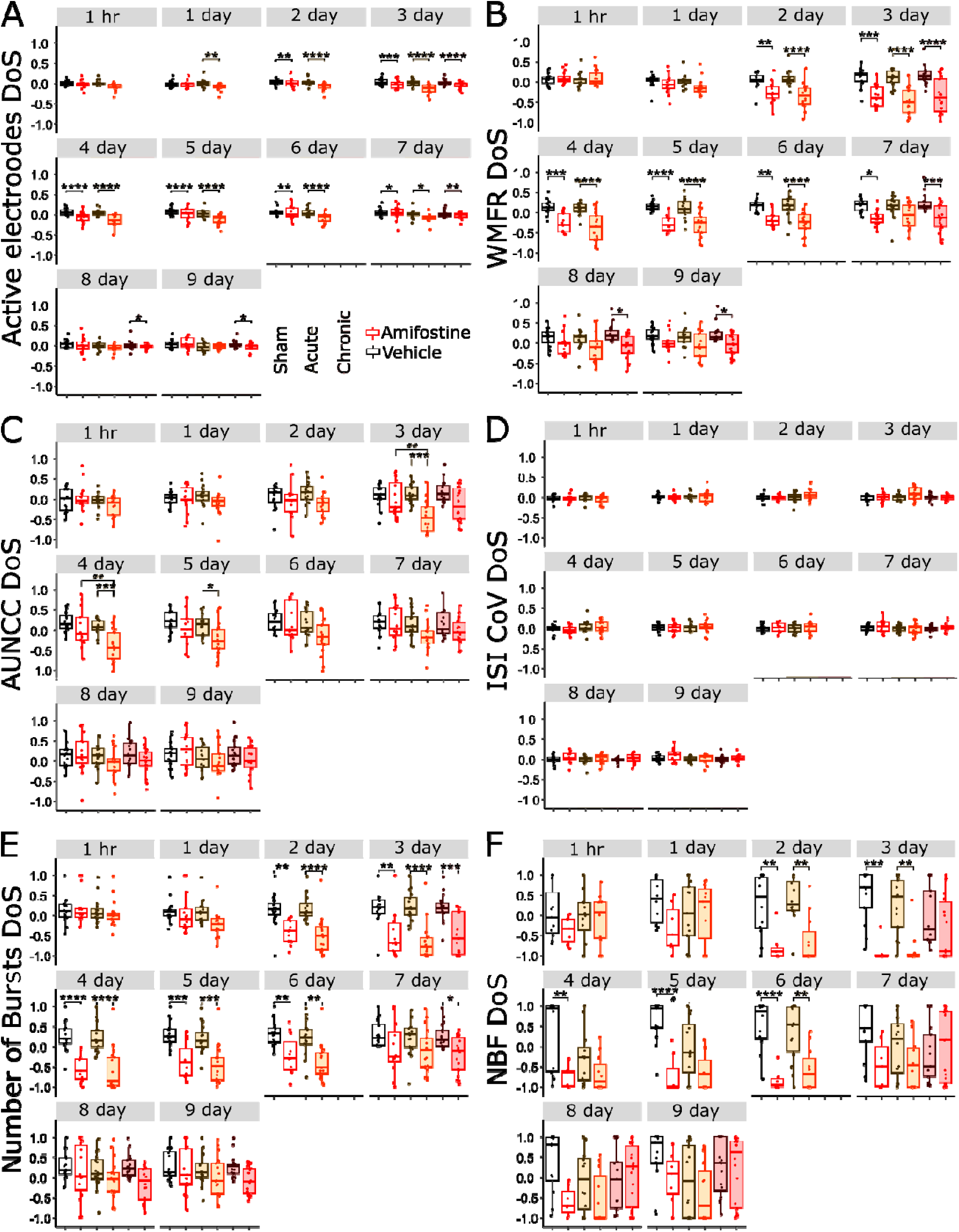
Chronic gamma radiation effects on electrophysiology. 2D mECN cultures treated with sham (white background) acute (yellow background) or chronic (red background) gamma radiation without (vehicle, black outline) or with (red outline) amifostine incubation. Enumerated panels visualize DoS of A. AE, B. WMFR, C. AUNCC, D. ISI CoV, E. number of bursts, and F. NBF. Significant effects due to amifostine indicated with * while significant effects of radiation indicated with #.

## Discussion

The detrimental effects of IR on CNS function is a major concern for missions venturing beyond low earth orbit, including those to the lunar surface, lunar orbit, or the Martian surface (Patel, Brunstetter et al. 2020, Kerry O’Banion 2022, Seidler, Mao et al. 2024). The importance of understanding such effects cannot be understated as this health risk carries a high likelihood of occurrence and high adverse consequences (Patel, Brunstetter et al. 2020). Data on the long-term effects of chronic IR exposure, especially that which is relevant to space radiation exposure, is highly limited and, as such, our understanding of the cell and tissue level effects of chronic IR exposure is not well characterized (Chancellor, Blue et al. 2018, Cao, Weil et al. 2022). The application of *in vitro* 3D CNS models, as described here, provides the unique opportunity to isolate neuroglial cell- and tissue-level responses to acute and chronic IR. Parsing the effects of acute and chronic space radiation relevant doses on CNS tissue function will be essential for understanding the biological effects that may not be fully captured in studies that utilize only acute doses.

This study utilizes an engineered 3D *in vitro* type-I collagen-based CTM in combination with 2D cultures seeded on MEA plates to understand the phenotypic changes in functional readouts of acute and chronic IR exposure, including neuroimmune reactivity and network level electrophysiology (Lojek, Bostanci et al. 2026) (Lojek, Williams et al. 2024). The inclusion of the FDA-approved radioprotectant drug amifostine (King, Joseph et al. 2020), known to reduce DNA damage due to IR, is intended to demonstrate proof of concept for drug screening using this platform. The DNA damage response from acute IR exposure is among the most well characterized genomic injury phenotypes (Mavragani, Nikitaki et al. 2019). Assessment of DNA damage was included as a validation step in our study and primarily aimed to verify the presence of a well characterized biological response to IR exposure prior to the collection of functional injury readouts (Figure 2B,E). The use of DNA damage markers such as 53BP1(Schultz, Chehab et al. 2000) is commonplace in radiation biology and space radiation studies, as DNA damage repair remains one of few reliable IR injury markers. 53BP1 is most often measured at early timepoints (≤ 24 h) and while many studies count radiation induced foci in cell nuclei at high magnification, 53BP1 nuclear intensity has also been used as a measure of DNA damage(Furia, Pelicci et al. 2022). Our results show an increase in the intensity of 53BP1, a DNA damage repair protein, at 30 min post-acute IR exposure as is consistent with published finding (Lojek, Bostanci et al. 2026) (Figure 2A,B). This phenotype is mitigated in the amifostine-treated acute exposure samples at 30 min post-IR in agreement with our previous work (Lojek, Williams et al. 2024) indicating that amifostine mitigates against acute gamma radiation-induced DNA damage (Figure 2E). In chronically treated samples, at 3-days of exposure (cumulative 0.200 Gy), both vehicle- and amifostine-treated samples experience a significant decrease in the intensity of nuclear 53BP1 relative to sham (Figure 2 C,F). Interestingly, at 7 days (0.47 Gy cumulative) both IR exposed vehicle- and amifostine-treated samples exhibit a significant increase in 53BP1 intensity relative to controls (Figure 2 D,G). These data suggest there may be fluctuations in the ratios of proliferative to non-proliferative cells over time or temporary exhaustion of DNA repair capacity (Wheeler and Nelson 1991) {Ahmed, 1979 #71} though this requires further study to adequately describe (Figure 2D,G). Briefly, 53BP1 dynamics are not well characterized at protracted timepoints (beyond 24 h) in cases of chronic exposure (Osipov, Chigasova et al. 2024) and there could be dynamic and yet uncharacterized mechanisms in play at protracted timepoints (>7 days) due to ROS accumulation or constitutively activated DNA damage repair mechanisms. Overall, these results confirmed DNA damage in CTM samples 30 min after acute irradiation indicating that cells are responding to IR exposure in a manner consistent with existing literature(Schultz, Chehab et al. 2000) (Osipov, Chigasova et al. 2024).

Immune reactivity in the CNS is characterized by the upregulation of pro-inflammatory proteins associated with a pro-inflammatory response (Sochocka, Diniz et al. 2017). CNS cell-type specific immune responses chiefly involve microglia, the resident macrophages of the brain(Kerry O’Banion 2022, Dolan, Therrien et al. 2023), and astrocytes, glial cells essential in supporting and regulating synapses (Zhao, Huang et al. 2024). In acute but not chronic gamma irradiated samples, CTM GFAP expression increased irrespective of amifostine treatment (Figure 3C), indicating an acute immune response. This indicates the presence of a mild immune response to acute radiation but not chronic (9 days) exposure. GFAP cell object circularity is significantly increased in acute and chronic irradiated CTMs without amifostine (Figure 3D). The lack of a pro-inflammatory circularity phenotype in chronic-amifostine treated samples could indicate that amifostine confers protection against long-term injury in astrocytes, suggesting amifostine may be more effective at preventing accumulation of ROS over time than mitigating acute DNA damage at lower dose rates. It is also possible that chronic radiation exposure elicits a delayed neuroimmune response (> 9 days) in the CTM model due to accumulating DNA damage. Such a response would be consistent with *in vivo* murine literature which shows inflammatory responses occurring on the order of months (Prezado, Jouvion et al. 2017) (Tang and et al. 2022) and would require longitudinal study which is beyond the scope of the current work. The observed cytomorphology changes, including a reduction of astrocyte circularity phenotype, is typically indicative of an activated state (Hyvärinen, Hagman et al. 2019, Delgado-García, Ojalvo-Sanz et al. 2024) (Figure 3D). Despite the increased astrocyte circularity, no significant difference is found in either perimeter or Feret diameter (Figure E, F), indicating that slight cytoskeletal changes occur due to both acute and chronic radiation and appear correlated GFAP intensity change. In summary our data shows that astrocyte reactivity is increased albeit without characteristic responses expected from a major injury such as endotoxin exposure (Diaz-Castro, Bernstein et al. 2021, Sun, Song et al. 2025).

We observed no change in the number of NeuN+ neurons across conditions at the terminal timepoint (9 days following the start of IR or sham exposure) indicating that neuronal cell number is not affected by acute or chronic gamma radiation within our timeline. This NeuN result agrees with our previous findings at slightly earlier timepoints (up to 7 days) (Lojek, Bostanci et al. 2026). Overall, these findings indicate a mild inflammation mediated by astrocytes, which is in agreement with other CNS irradiation studies (Verma, Passerat de la Chapelle et al. 2022, Oyefeso, Goldberg et al. 2023) although it should be noted that, in some cases, even higher IR doses (2 Gy) had no effect on astrocyte morphology in human brain organoids at 24 h following exposure (Oyefeso, Goldberg et al. 2023). Importantly, amifostine does not appear to convey a protective effect against chronic astrocyte reactivity. The authors are not aware of any existing literature examining the effects of amifostine on astrocytes.

Iba1 is a macrophage marker closely associated with microglia activation state and is commonly used to identify presumptive microglia (Ohsawa, Imai et al. 2004, Lier, Streit et al. 2021). We observed that both acute and chronic gamma radiation reduced Iba1 expression in CTMs (Figure 4 A, C). While Iba1+ cell-object circularity is significantly altered in chronic-amifostine samples relative to sham-amifostine controls (Figure 4 D), Iba1 cell object perimeter (Figure 4 E) and Feret diameter (Figure 4F) show significant decreases in acute-vehicle and chronic-vehicle samples relative to sham-vehicle controls. These data suggest that gamma irradiation significantly impacts microglial cell structure, perhaps triggering phenotypic switching from a ramified to ameboid morphology indicative of immune activation (Lier, Streit et al. 2021).

Typically, the immune response to radiotherapy-relevant acute doses (between 10 and 25 Gy), has been reported to increase Iba1 expression in irradiated mouse brain between protracted timepoints of 6 to 12 months (Prezado, Jouvion et al. 2017) (Tang and et al. 2022). On the other hand, the murine microglia cell line BV2 exhibits a decrease in the expression of some pro-inflammatory RNA transcripts, when treated with a comparable 1 Gy X-ray dose (Kim, Chung et al. 2020), which is consistent with the partial anti-inflammatory response in our ICC results. Similarly, a study using the human microglial cell line CHME 5 shows that immune reactivity is highly dependent on cumulative dose (0.5, 1, 2, 4, 8 Gy) and temporal sampling (2 h to 14 days) (Chen, Chong et al. 2016). Thus, the reduction in Iba1 that has been observed in our model differs from a whole-organism inflammatory response observed in animals but falls in line with the limited *in vitro* literature. Our model may reflect a snapshot (9 days following the start of exposure) of a single point in a complex response that changes over time, and supports the need for further exploration in a longitudinal study dedicated to exploring long-term immune responses. Similarly to its effects on astrocyte activation, amifostine also reduces microglial reactivity in both sham and irradiated samples based on Iba1 upregulation and cytomorphology (Figure 4C,E,F). There have been no studies specifically examining Iba1 expression changes in microglia after amifostine treatment. Together these results indicate that a limited anti-inflammatory immune response due to both acute and chronic radiation is occurring in both astrocytes and microglia, with radioprotective effects of amifostine observed in microglia but not astrocyteswhich is consistent with our previous findings (Lojek, Bostanci et al. 2026).

To further assess the immune response induced by acute and chronic radiation, media samples were retained from both CTM and 2D MEA cultures prior to experimentation and at 7-days following the beginning of chronic (or the administration of acute) IR exposure. There is no substantial increase in the pro-inflammatory markers IL-1β, KC, MCP-1, or MIP-2 in either the 2D or CTM samples to indicate the presence of a pro-inflammatory immune response (Figure 5A, B). We report a significant relative decrease in vehicle-chronic IL-6 and in amifostine-chronic KC, MCP1, and MIP2 relative to sham-vehicle control for CTM samples. We assess that these decreases are not indicative of a noteworthy anti-inflammatory response as relatively small fluctuations in these pro-inflammatory cytokines are not necessarily indicative of a phenotypic change. While Verma et al. (2022) (Verma, Passerat de la Chapelle et al. 2022) does document an increase in pro-inflammatory cytokine secretions in astrocyte and endothelial cell cultures this platform is fundamentally different from that which we describe because of cell types included (neurons, astrocytes, and microglia), and the hydrogel matrix compared to Matrigel or collagen. Additionally, the Verma et al. (2022) platform includes endothelial cells which could be driving cytokine production. Conversely, a BV-2 murine microglia based platform has demonstrated an anti-inflammatory effect of IR, where pro-inflammatory cytokines TNF-α (below the detection limit in the present study), IL-1β, and IL-6 (below the detection limit in the present study) were all decreased in irradiated samples when stimulated with lipopolysaccharide(Kim, Chung et al. 2020). This further supports our findings indicating that gamma radiation may suppress an inflammatory phenotype in CNS, however further study would be necessary to confirm this. Overall these results in combination with the observed changes in Iba1 and GFAP indicate that changes to inflammatory state are subtle in this model. Naturally, this raises further questions about the nature of radiation induced neuroimmune phenotypes and their role in behavioral changes. A future study specifically examining the effects of IR, ideally using a broad dose curve and incorporating transcriptomic analysis, would provide valuable information on the immune response of microglia.

A significant component of IR injury is believed to occur through the generation of free radical species in the cell lumen. Free radicals react with biomolecules leading to cellular dysfunction, such as failure to pass DNA-damage checkpoints (Iliakis, Wang et al. 2003) or the expression of oncogenes (Garte and Burns 1991) (Kumar, Kumar et al. 2023). The IR injury process can be mimicked by “radiomimetic” drugs that generate free radicals within the cell, leading to DNA damage. Bleomycin is a radiomimetic enediyne routinely used as a positive control for oxidative stress and DNA damage. (Povirk 1996) (Shimizu, Izawa et al. 2025).

We treated 2D mECN cultures grown on MEAs with bleomycin to identify the effects that a radiomimetic reactive oxygen species-induced injury has on network-level electrophysiology and compare it to electrophysiology changes induced by acute and chronic gamma radiation. Additionally, the use of bleomycin allowed us to map the dose response of the free-radical mediated DNA damage beyond gamma radiation levels used in the experiment. Our results show that the number of active electrodes remains essentially unchanged at 1, 24, and 48 h following treatment except at the highest bleomycin concentration (50 µg/mL) indicating that widespread cell death is likely not occurring at lower concentrations (0.01 to 10 µg/mL) (Figure 6F-K). Based on existing literature a bleomycin dose of 1 µg/mL is equivalent to an acute IR dose of approximately 0.4 Gy (Mladenov, Kalev et al. 2009, Lu, Zhang et al. 2017), however the authors could find no empirical information to support this equivalence. Reductions in WMFR at 1 h and 48 h as well as in AUNCC, number of bursts, and NBF at 48 h from high bleomycin doses (50 µg/mL) but not at lower concentrations (0.01 to 10 µg/mL) indicates electrophysiological activity seems to be largely unaffected by bleomycin. This suggests that mECN are highly robust to ionizing radiation-like injury even at doses potentially in excess of approximately a 4 Gy equivalent injury (Mladenov, Kalev et al. 2009, Lu, Zhang et al. 2017).

Electrophysiology recordings were made on 2D mECN cultures prior to gamma radiation exposure or amifostine treatment, at 1 h, and daily for 9 days following the beginning of gamma radiation exposure. Our results show little or no change in MEA metrics including number of active electrodes, firing rate (WMFR), synchrony (AUNCC), regularity (ISI CoV), bursting, or network bursting (NBF) due to either acute radiation or chronic radiation alone (Figure 7 A-F). These results suggest that 2D mECN electrophysiological phenotype is not directly affected by a 0.5 Gy dose of gamma radiation via acute or chronic exposure. These findings are in keeping with our previous results which indicate that proton radiation (10 Gy) does not alter electrophysiology activity (Lojek, Bostanci et al. 2026). Importantly, the MEA analysis techniques used here are based on descriptive statistics derived from whole (10 min) recordings, and while these techniques are well established, they could be missing subtle phenotypes related to complex network activity in the data such as “reverberation” patterns within bursts (Hernandes, Heuvelmans et al. 2024). For this reason, future analysis incorporating advanced statistical techniques might yield subtle phenotypes undetected by traditional analysis.

Amifostine exerts electrophysiological effects on these cultures, significantly reducing WMFR and number of bursts from days 2 through 6 of exposure. Despite the decrease in WMFR, there is minimal evidence that amifostine causes cell death, as the number of active electrodes is not significantly reduced (Figure 7A). This agrees with our previous work which shows minimal amifostine cytotoxicity (Lojek, Bostanci et al. 2026). The electrophysiological effect of amifostine found here could explain the well documented CNS side effects of this drug which include neurological symptoms such as nausea and vomiting (Rades, Fehlauer et al. 2004).

In conclusion, space-relevant doses of acute and chronic gamma radiation evoke contradictory effects on inflammation in a 3D CNS model, with no changes in neuronal morphology and function. Specifically, astrocytes adopt a reactive phenotype while microglial shows altered activation state. Neither of these effects are strictly regulated by radiation dose rate. Further, these findings suggest that neuronal network function is highly resistant to both acute and chronic gamma radiation at the doses and dose rates used in this study. We incidentally report that amifostine modulates neuronal network function, which could explain documented CNS associated side effects of the drug (Rades, Fehlauer et al. 2004), and has an anti-inflammatory effect on microglia. Together these results suggest that mitigation approaches to chronic low-dose ionizing radiation may focus on glial dysfunction rather than neuronal health, and that the neuronal effects of amifostine limit its use as a spaceflight countermeasure. Further studies utilizing this model to define the CNS risk during deep space exploration may include exposures to real or simulated galactic cosmic radiation and comparison between mouse and human CTM radiosensitivity. This work has several limitations that should be considered. Here, we use sparsely ionizing gamma radiation to emulate injury caused by galactic cosmic radiation which is mostly composed of densely ionizing particles. Additionally, chronic exposure here is delivered over only 7 days whereas crewed missions beyond low earth orbit will experience IR exposure for longer durations (months) and at lower dose rates. Future studies to address this important limitation would likely require CNS-specific tissue culture models capable of remote reporting to be incorporated as payloads on uncrewed deep space research missions.

## Acknowledgements

The authors are thankful to Grace Callen for technical support on the Luminex Assay.

## Author Contributions

Neal M. Lojek, Chiara E. Ghezzi, Bryan J. Black, Andrew Rogers, Egle Cekanaviciute, and Erno Sajo contributed to study design. Neal M. Lojek, Nazli Bostanci, Paulo Henrique Borges, Chiara E. Ghezzi, and Bryan J. Black contributed to cell culture, construct generation, data collection, and data analysis. Andrew M. Rogers, Neal M. Lojek, and Nazli Bostanci contributed to culture irradiations.

## Data availability statement

Data generated in the making of this manuscript can be provided upon reasonable request.

