## Supplementary material for "Electrophysiological and neuroimmune responses of a cortical tissue model to chronic gamma radiation exposure": Supp Figures and Tables

**Supplemental information**

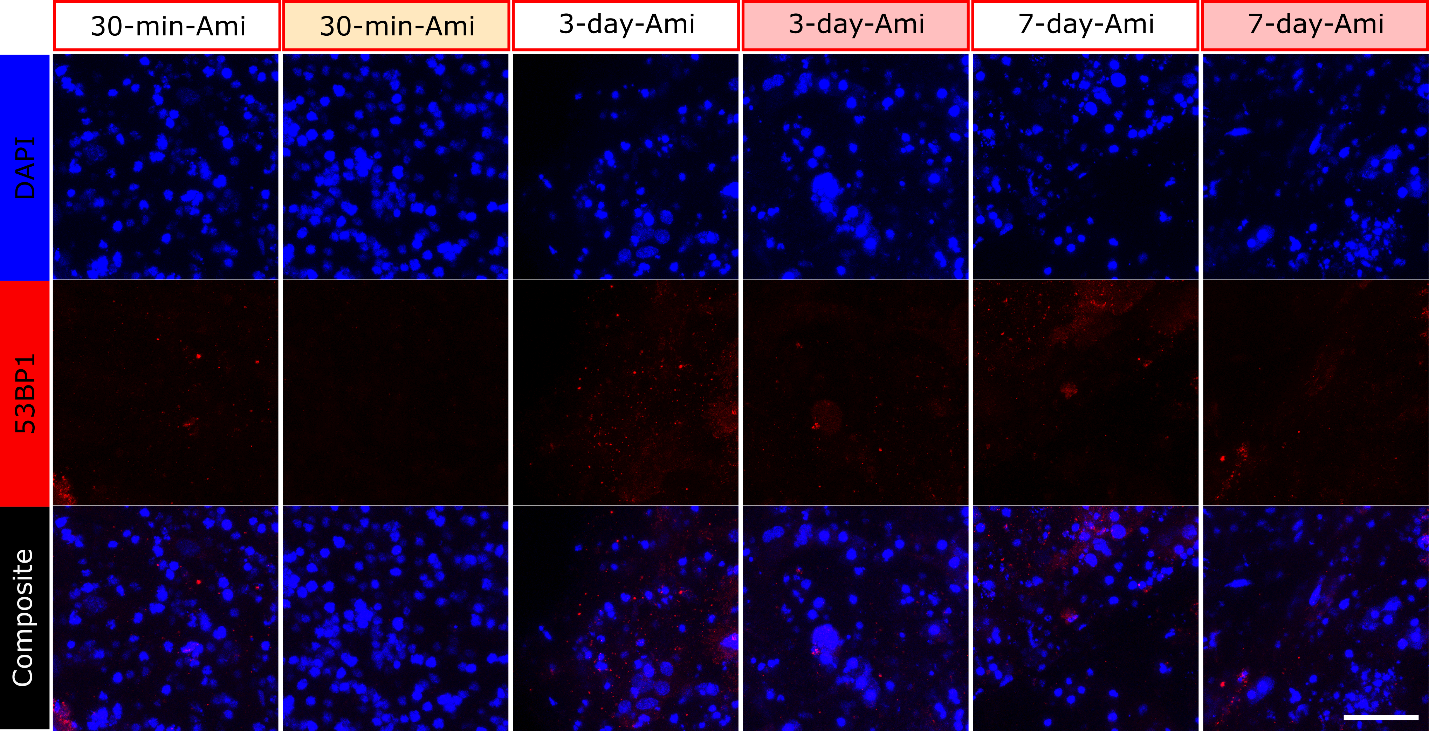

Figure S1. Representative CLSM images of DAPI (blue) and 53BP1 (red) ICC staining in sham (white) acute (light yellow) and chronic (light red) cultures pre-treated with Amifostine. Scale bar = 50 µm.

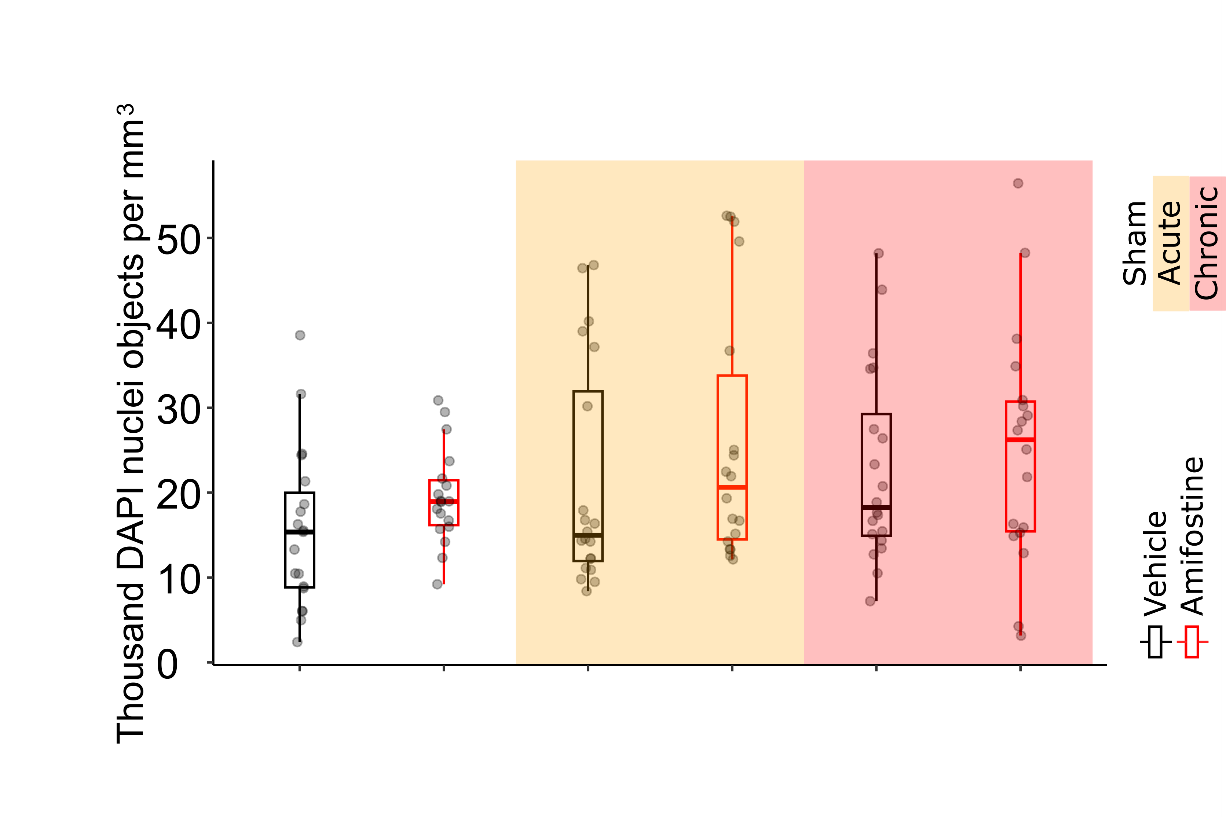

Figure S2. DAPI nuclei per mm3 as measured for Sham (unshaded), acute (yellow shaded) or chronic (pink shaded) radiation treatment and vehicle (black outline) and amifostine (red outline) pre-treatment at the 9-day timepoint.

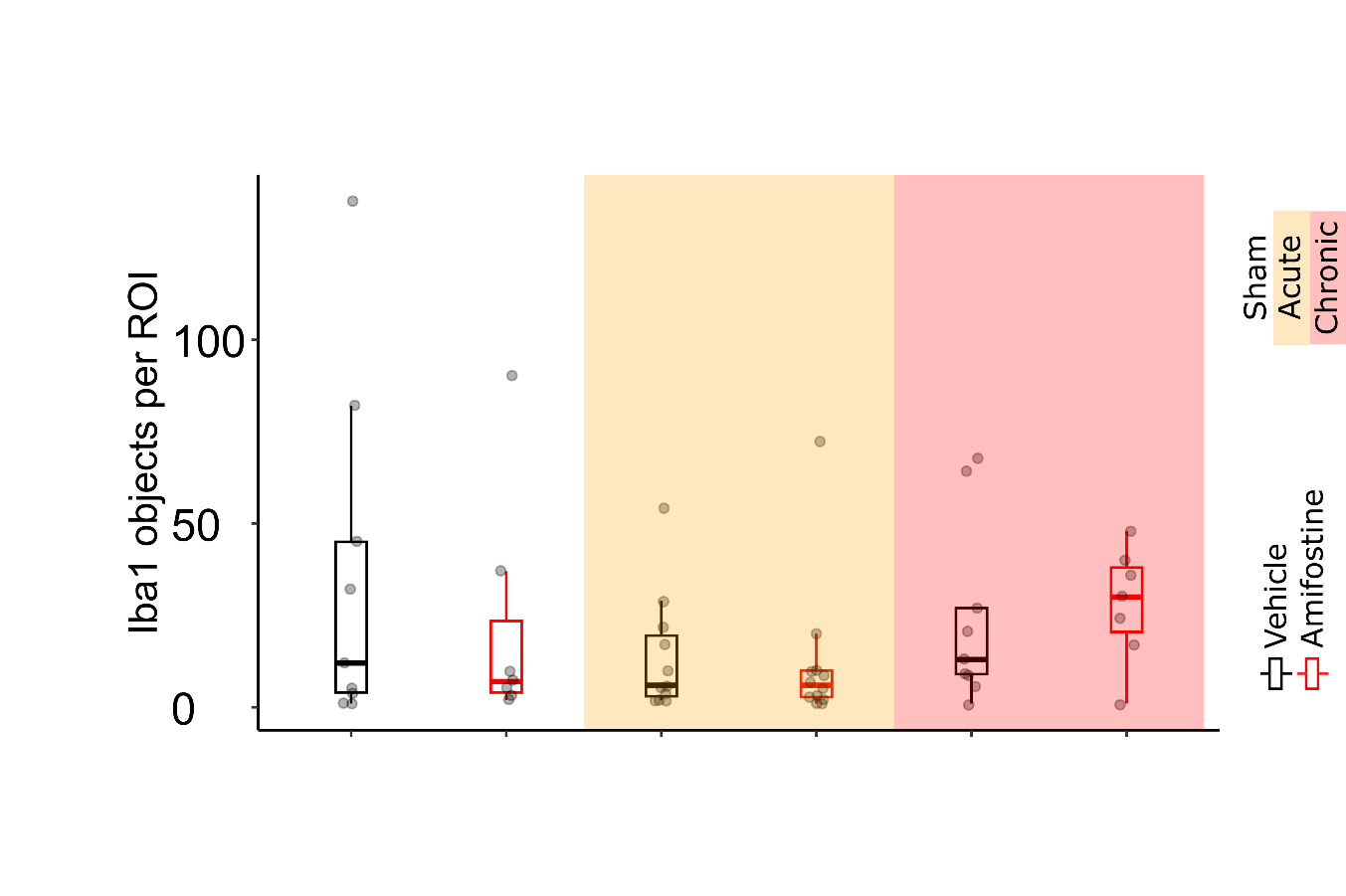

Figure S3. Iba1 cell number measured for Sham (unshaded), acute (yellow shaded) or chronic (pink shaded) radiation treatment and vehicle (black outline) and amifostine (red outline) pre-treatment at the 9-day timepoint.

|  | group1 | group2 | p.adj | p.adj.signif | GFAP_metric |
| --- | --- | --- | --- | --- | --- |
| 1 | Chronic-Amifostine | Sham-Vehicle | 0.0405 | * | Area |
| 2 | Acute-Amifostine | Chronic-Amifostine | 4.97e-06 | **** | Circ. |
| 3 | Acute-Amifostine | Sham-Amifostine | 3.82e-06 | **** | Circ. |
| 4 | Acute-Amifostine | Sham-Vehicle | 3.21e-05 | **** | Circ. |
| 5 | Acute-Vehicle | Chronic-Amifostine | 8.09e-08 | **** | Circ. |
| 6 | Acute-Vehicle | Chronic-Vehicle | 0.0362 | * | Circ. |
| 7 | Acute-Vehicle | Sham-Amifostine | 5.82e-08 | **** | Circ. |
| 8 | Acute-Vehicle | Sham-Vehicle | 9.23e-07 | **** | Circ. |
| 9 | Chronic-Amifostine | Chronic-Vehicle | 0.0075 | ** | Circ. |
| 10 | Chronic-Vehicle | Sham-Amifostine | 0.00658 | ** | Circ. |
| 11 | Chronic-Vehicle | Sham-Vehicle | 0.0177 | * | Circ. |
| 12 | Acute-Amifostine | Chronic-Amifostine | 4.4e-07 | **** | Mean |
| 13 | Acute-Amifostine | Sham-Amifostine | 4.3e-08 | **** | Mean |
| 14 | Acute-Amifostine | Sham-Vehicle | 0.00237 | ** | Mean |
| 15 | Acute-Vehicle | Chronic-Amifostine | 5.06e-14 | **** | Mean |
| 16 | Acute-Vehicle | Chronic-Vehicle | 9.11e-06 | **** | Mean |
| 17 | Acute-Vehicle | Sham-Amifostine | 1.78e-15 | **** | Mean |
| 18 | Acute-Vehicle | Sham-Vehicle | 4.78e-08 | **** | Mean |
| 19 | Chronic-Amifostine | Chronic-Vehicle | 0.00963 | ** | Mean |
| 20 | Chronic-Vehicle | Sham-Amifostine | 0.00237 | ** | Mean |

Supplemental table 1. All GFAP significant reactivity and cytomorphology pairwise comparisons by Holm-Bonferroni test.

|  | group1 | group2 | p.adj | p.adj.signif | Iba1_metric |
| --- | --- | --- | --- | --- | --- |
| 1 | Acute-Vehicle | Sham-Vehicle | 0.00942 | ** | Area |
| 2 | Chronic-Amifostine | Sham-Vehicle | 0.00014 | *** | Area |
| 3 | Chronic-Vehicle | Sham-Vehicle | 0.0134 | * | Area |
| 4 | Sham-Amifostine | Sham-Vehicle | 0.00556 | ** | Area |
| 5 | Acute-Vehicle | Chronic-Amifostine | 0.0271 | * | Circ |
| 6 | Chronic-Amifostine | Sham-Amifostine | 0.0314 | * | Circ |
| 7 | Acute-Amifostine | Sham-Vehicle | 0.0421 | * | Feret |
| 8 | Acute-Vehicle | Sham-Vehicle | 0.00632 | ** | Feret |
| 9 | Chronic-Amifostine | Chronic-Vehicle | 0.0422 | * | Feret |
| 10 | Chronic-Amifostine | Sham-Vehicle | 1.51E-08 | **** | Feret |
| 11 | Chronic-Vehicle | Sham-Vehicle | 0.0203 | * | Feret |
| 12 | Sham-Amifostine | Sham-Vehicle | 0.00103 | ** | Feret |
| 13 | Acute-Amifostine | Chronic-Vehicle | 0.0329 | * | Mean |
| 14 | Acute-Amifostine | Sham-Vehicle | 1.73E-08 | **** | Mean |
| 15 | Acute-Vehicle | Sham-Vehicle | 0.000263 | *** | Mean |
| 16 | Chronic-Amifostine | Sham-Vehicle | 3.49E-05 | **** | Mean |
| 17 | Chronic-Vehicle | Sham-Vehicle | 0.00714 | ** | Mean |
| 18 | Sham-Amifostine | Sham-Vehicle | 2.99E-05 | **** | Mean |
| 19 | Acute-Vehicle | Sham-Vehicle | 0.0159 | * | Perim |
| 20 | Chronic-Amifostine | Sham-Vehicle | 9.11E-06 | **** | Perim |
| 21 | Chronic-Vehicle | Sham-Vehicle | 0.0245 | * | Perim |
| 22 | Sham-Amifostine | Sham-Vehicle | 0.0148 | * | Perim |

Supplemental table 2. All significant Iba1 reactivity and cytomorphology pairwise comparisons by Holm-Bonferroni test.

| drug_status | time | mean_AE | sd_AE | mean_WMFR | SD_WMFR | mean_AUNCC | sd_AUNCC | mean_ISI_CoV | sd_ISI_CoV | mean_NumB | sd_NumB | mean_NBF | sd_NBF |
| --- | --- | --- | --- | --- | --- | --- | --- | --- | --- | --- | --- | --- | --- |
| 0.01 µg/mL | 1 hour | 0.0000 | 0.0000 | -0.0203 | 0.0000 | 0.0093 | 0.1307 | -0.0560 | 0.0875 | -0.0151 | 0.1007 | -0.1311 | 0.4693 |
| 0.01 µg/mL | 24 hour | 0.0179 | 0.0357 | 0.0256 | 0.0357 | 0.0782 | 0.1699 | -0.0043 | 0.1256 | 0.0428 | 0.1917 | -0.4285 | 0.3030 |
| 0.01 µg/mL | 48 hour | 0.0006 | 0.0292 | 0.1018 | 0.0292 | 0.0627 | 0.1866 | -0.0220 | 0.1067 | 0.1226 | 0.2445 | -0.1361 | 0.6865 |
| 0.1 µg/mL | 1 hour | 0.0172 | 0.0199 | 0.0045 | 0.0199 | -0.0377 | 0.0275 | -0.0087 | 0.0045 | -0.0742 | 0.0931 | 0.0112 | 0.5949 |
| 0.1 µg/mL | 24 hour | 0.0000 | 0.0000 | 0.0601 | 0.0000 | 0.0621 | 0.1340 | -0.0061 | 0.0289 | 0.0314 | 0.1622 | -0.1706 | 0.6906 |
| 0.1 µg/mL | 48 hour | 0.0345 | 0.0488 | 0.0294 | 0.0488 | 0.0990 | 0.0715 | 0.0024 | 0.0213 | -0.0133 | 0.2709 | -0.4336 | 0.5280 |
| 1 µg/mL | 1 hour | -0.0208 | 0.0417 | 0.0623 | 0.0417 | 0.0259 | 0.0504 | 0.0198 | 0.0453 | 0.0445 | 0.0382 | 0.1450 | 0.2652 |
| 1 µg/mL | 24 hour | -0.0100 | 0.0200 | 0.0840 | 0.0200 | 0.0642 | 0.0641 | 0.0117 | 0.0458 | 0.0822 | 0.0816 | 0.5461 | 0.4830 |
| 1 µg/mL | 48 hour | -0.0228 | 0.0500 | 0.2029 | 0.0500 | 0.1179 | 0.1948 | -0.0025 | 0.0413 | 0.2340 | 0.1420 | 0.6724 | 0.5175 |
| 10 µg/mL | 1 hour | 0.0000 | 0.0000 | 0.0556 | 0.0000 | 0.0549 | 0.1594 | -0.0090 | 0.0309 | 0.0518 | 0.0267 | 0.2180 | 0.3332 |
| 10 µg/mL | 24 hour | -0.0189 | 0.0223 | 0.0988 | 0.0223 | -0.1080 | 0.3296 | -0.0157 | 0.0467 | 0.1177 | 0.1952 | 0.3448 | 0.4295 |
| 10 µg/mL | 48 hour | -0.0417 | 0.0833 | 0.0844 | 0.0833 | -0.0105 | 0.3478 | 0.0287 | 0.0849 | 0.1101 | 0.2107 | 0.4606 | 0.2215 |
| 50 µg/mL | 1 hour | -0.0145 | 0.0627 | -0.0510 | 0.0627 | 0.0112 | 0.0281 | -0.0192 | 0.0394 | 0.1895 | 0.5442 | -0.4533 | 0.5402 |
| 50 µg/mL | 24 hour | -0.1024 | 0.1208 | -0.1624 | 0.1208 | -0.3473 | 0.4323 | -0.0090 | 0.0443 | -0.0168 | 0.7168 | -0.7091 | 0.4860 |
| 50 µg/mL | 48 hour | -0.3043 | 0.2009 | -0.5569 | 0.2009 | -0.6561 | 0.3098 | 0.0166 | 0.0509 | -0.6297 | 0.3162 | -1.0000 | 0.0000 |
| Vehicle | 1 hour | -0.0045 | 0.0359 | 0.0373 | 0.0359 | 0.0008 | 0.0870 | -0.0576 | 0.0580 | 0.0180 | 0.0787 | -0.1640 | 0.5653 |
| Vehicle | 24 hour | 0.0007 | 0.0554 | 0.0560 | 0.0554 | 0.1066 | 0.1415 | -0.0271 | 0.0979 | 0.0739 | 0.1890 | 0.0709 | 0.0490 |
| Vehicle | 48 hour | 0.0573 | 0.0558 | 0.0619 | 0.0558 | 0.1112 | 0.1519 | -0.0416 | 0.0776 | 0.1726 | 0.1236 | 0.3582 | 0.1767 |

Supplemental table 3. Bleomycin treatment results for MEA metrics shown as mean and standard deviation (sd) DoS normalized.

| all_stat | time | mean_AE | sd_AE | mean_WMFR | SD_WMFR | mean_AUNCC | sd_AUNCC | mean_ISI_CoV | sd_ISI_CoV | mean_NumB | sd_NumB | mean_NBF | sd_NBF |
| --- | --- | --- | --- | --- | --- | --- | --- | --- | --- | --- | --- | --- | --- |
| Acute_amifostine | 1 hr | -0.0307 | 0.0407 | 0.0335 | 0.0407 | -0.0126 | 0.1215 | 0.0009 | 0.0522 | 0.0607 | 0.3210 | 0.1124 | 0.4862 |
| Acute_amifostine | 1 day | -0.0770 | 0.0624 | -0.1630 | 0.0624 | 0.0131 | 0.2056 | 0.0682 | 0.1224 | -0.1934 | 0.3612 | 0.3738 | 0.6061 |
| Acute_amifostine | 2 day | -0.0888 | 0.1004 | -0.4036 | 0.1004 | -0.0599 | 0.2914 | 0.0690 | 0.1189 | -0.5040 | 0.5479 | -0.4897 | 0.6663 |
| Acute_amifostine | 3 day | -0.1712 | 0.1429 | -0.5805 | 0.1429 | -0.4342 | 0.4147 | 0.1030 | 0.1710 | -0.7074 | 0.5300 | -0.8199 | 0.4868 |
| Acute_amifostine | 4 day | -0.1450 | 0.1495 | -0.4845 | 0.1495 | -0.3761 | 0.4492 | 0.0788 | 0.1569 | -0.6241 | 0.5852 | -0.6804 | 0.4886 |
| Acute_amifostine | 5 day | -0.1551 | 0.1256 | -0.4191 | 0.1256 | -0.2661 | 0.4298 | 0.0577 | 0.1428 | -0.4884 | 0.5354 | -0.6475 | 0.4714 |
| Acute_amifostine | 6 day | -0.1415 | 0.1104 | -0.3856 | 0.1104 | -0.2085 | 0.4973 | 0.0792 | 0.1308 | -0.4147 | 0.5405 | -0.5107 | 0.5929 |
| Acute_amifostine | 7 day | -0.1130 | 0.0922 | -0.2516 | 0.0922 | -0.2272 | 0.3898 | 0.0422 | 0.0841 | -0.2735 | 0.5771 | -0.6271 | 0.5197 |
| Acute_amifostine | 8 day | -0.0981 | 0.0864 | -0.2552 | 0.0864 | -0.1254 | 0.3835 | 0.0647 | 0.0821 | -0.1868 | 0.5500 | -0.5325 | 0.6837 |
| Acute_amifostine | 9 day | -0.0499 | 0.0529 | -0.2203 | 0.0529 | -0.1503 | 0.3941 | 0.0348 | 0.0879 | -0.1591 | 0.5614 | -0.6036 | 0.5653 |
| Acute_Vehicle | 1 hr | 0.0171 | 0.0829 | 0.0163 | 0.0829 | 0.0413 | 0.0727 | -0.0006 | 0.0548 | 0.0581 | 0.1367 | 0.0893 | 0.6119 |
| Acute_Vehicle | 1 day | 0.0171 | 0.0350 | -0.0270 | 0.0350 | 0.0313 | 0.1677 | 0.0032 | 0.0423 | 0.0212 | 0.1503 | 0.2554 | 0.7259 |
| Acute_Vehicle | 2 day | 0.0505 | 0.0742 | -0.0030 | 0.0742 | 0.0932 | 0.1490 | 0.0110 | 0.0974 | 0.1291 | 0.2472 | 0.3485 | 0.6769 |
| Acute_Vehicle | 3 day | 0.0286 | 0.0340 | 0.0182 | 0.0340 | 0.0100 | 0.1710 | -0.0197 | 0.0567 | 0.1255 | 0.2879 | 0.1331 | 0.8694 |
| Acute_Vehicle | 4 day | 0.0551 | 0.0748 | 0.0446 | 0.0748 | 0.0755 | 0.1217 | -0.0110 | 0.0511 | 0.1817 | 0.3053 | -0.3275 | 0.7231 |
| Acute_Vehicle | 5 day | 0.0373 | 0.0510 | 0.0355 | 0.0510 | 0.0077 | 0.2647 | -0.0368 | 0.0614 | 0.1246 | 0.3568 | 0.2031 | 0.7810 |
| Acute_Vehicle | 6 day | 0.0442 | 0.0917 | 0.0278 | 0.0917 | 0.0241 | 0.3014 | -0.0423 | 0.0815 | 0.0957 | 0.4682 | 0.3044 | 0.7878 |
| Acute_Vehicle | 7 day | 0.0160 | 0.0633 | 0.0644 | 0.0633 | 0.0097 | 0.2887 | -0.0367 | 0.0661 | 0.1261 | 0.5149 | 0.2498 | 0.7454 |
| Acute_Vehicle | 8 day | 0.0155 | 0.1205 | 0.0043 | 0.1205 | 0.0286 | 0.3549 | -0.0032 | 0.0610 | 0.1007 | 0.5039 | 0.2641 | 0.7218 |
| Acute_Vehicle | 9 day | -0.0267 | 0.0760 | 0.0639 | 0.0760 | 0.0028 | 0.3166 | -0.0103 | 0.0756 | 0.0998 | 0.3683 | 0.1209 | 0.8388 |
| Chronic_amifostine | 3 day | -0.0368 | 0.0825 | -0.3406 | 0.0825 | -0.1318 | 0.4183 | 0.0017 | 0.0677 | -0.4678 | 0.4991 | -0.3402 | 0.7914 |
| Chronic_amifostine | 7 day | -0.0190 | 0.0758 | -0.1432 | 0.0758 | -0.0255 | 0.3076 | 0.0318 | 0.0951 | -0.2083 | 0.4567 | 0.0190 | 0.8586 |
| Chronic_amifostine | 8 day | -0.0195 | 0.0545 | -0.0945 | 0.0545 | -0.0029 | 0.3341 | 0.0228 | 0.1138 | -0.1661 | 0.4161 | 0.0695 | 0.7494 |
| Chronic_amifostine | 9 day | -0.0237 | 0.0852 | -0.0332 | 0.0852 | 0.0300 | 0.3382 | 0.0312 | 0.0797 | -0.0689 | 0.3302 | 0.1953 | 0.8277 |
| Chronic_Vehicle | 3 day | 0.0371 | 0.0740 | 0.2323 | 0.0740 | 0.1783 | 0.2796 | 0.0103 | 0.0744 | 0.2782 | 0.3043 | 0.0574 | 0.8282 |
| Chronic_Vehicle | 7 day | 0.0306 | 0.0793 | 0.2885 | 0.0793 | 0.1967 | 0.3439 | 0.0043 | 0.0859 | 0.3020 | 0.3358 | -0.1774 | 0.8196 |
| Chronic_Vehicle | 8 day | 0.0288 | 0.1367 | 0.2740 | 0.1367 | 0.1825 | 0.3783 | -0.0218 | 0.0678 | 0.3086 | 0.3180 | 0.1259 | 0.7792 |
| Chronic_Vehicle | 9 day | 0.0342 | 0.1025 | 0.2687 | 0.1025 | 0.2374 | 0.3399 | -0.0062 | 0.0696 | 0.2962 | 0.3352 | 0.2627 | 0.6643 |
| Sham_amifostine | 1 hr | -0.0432 | 0.0860 | 0.0454 | 0.0860 | 0.0581 | 0.2764 | -0.0550 | 0.0588 | 0.0006 | 0.1246 | -0.3115 | 0.3212 |
| Sham_amifostine | 1 day | -0.0491 | 0.0484 | -0.0933 | 0.0484 | -0.0429 | 0.1545 | 0.0072 | 0.0550 | -0.1451 | 0.1963 | -0.1996 | 0.5514 |
| Sham_amifostine | 2 day | -0.0481 | 0.0469 | -0.2765 | 0.0469 | -0.0018 | 0.3679 | -0.0244 | 0.0481 | -0.4072 | 0.3055 | -0.7438 | 0.4105 |
| Sham_amifostine | 3 day | -0.0720 | 0.0625 | -0.5488 | 0.0625 | -0.1700 | 0.3843 | -0.0007 | 0.0988 | -0.7501 | 0.2054 | -0.8756 | 0.2926 |
| Sham_amifostine | 4 day | -0.1222 | 0.1137 | -0.4162 | 0.1137 | -0.0205 | 0.4900 | -0.0409 | 0.1073 | -0.7184 | 0.2040 | -0.6882 | 0.2588 |
| Sham_amifostine | 5 day | -0.0471 | 0.1075 | -0.3805 | 0.1075 | -0.0663 | 0.3525 | -0.0320 | 0.1133 | -0.5964 | 0.2243 | -0.6610 | 0.5211 |
| Sham_amifostine | 6 day | -0.0613 | 0.0743 | -0.2794 | 0.0743 | 0.0225 | 0.3977 | -0.0274 | 0.1107 | -0.4946 | 0.3199 | -0.8022 | 0.2786 |
| Sham_amifostine | 7 day | -0.0588 | 0.1482 | -0.2410 | 0.1482 | -0.0325 | 0.2895 | -0.0055 | 0.1225 | -0.4338 | 0.3559 | -0.4389 | 0.6404 |
| Sham_amifostine | 8 day | -0.0970 | 0.1144 | -0.1394 | 0.1144 | 0.0111 | 0.4820 | 0.0206 | 0.1617 | -0.3382 | 0.4210 | -0.5818 | 0.3939 |
| Sham_amifostine | 9 day | -0.0436 | 0.0785 | -0.1060 | 0.0785 | 0.0249 | 0.3309 | 0.0592 | 0.1508 | -0.2044 | 0.3343 | -0.1580 | 0.5600 |
| Sham_Vehicle | 1 hr | 0.0167 | 0.0463 | 0.0795 | 0.0463 | -0.0594 | 0.3034 | -0.0165 | 0.1028 | 0.0826 | 0.2820 | 0.1476 | 0.6935 |
| Sham_Vehicle | 1 day | -0.0606 | 0.0409 | 0.1040 | 0.0409 | 0.1217 | 0.1381 | 0.0493 | 0.0470 | 0.0796 | 0.0577 | 0.2623 | 0.3790 |
| Sham_Vehicle | 2 day | 0.0266 | 0.0275 | 0.1032 | 0.0275 | 0.1142 | 0.1580 | 0.0004 | 0.0897 | 0.1492 | 0.1881 | 0.1634 | 0.7760 |
| Sham_Vehicle | 3 day | -0.0022 | 0.0547 | 0.1520 | 0.0547 | 0.1356 | 0.1968 | 0.0053 | 0.0563 | 0.1537 | 0.1568 | 0.3951 | 0.6625 |
| Sham_Vehicle | 4 day | 0.0565 | 0.0551 | 0.1105 | 0.0551 | 0.2270 | 0.2309 | -0.0426 | 0.1333 | 0.1595 | 0.2122 | 0.2389 | 0.9058 |
| Sham_Vehicle | 5 day | 0.0266 | 0.0579 | 0.1168 | 0.0579 | 0.1412 | 0.3126 | -0.0026 | 0.0694 | 0.1207 | 0.2135 | 0.5934 | 0.3938 |
| Sham_Vehicle | 6 day | 0.0494 | 0.0341 | 0.1348 | 0.0341 | 0.1428 | 0.2878 | -0.0113 | 0.0833 | 0.1872 | 0.2100 | 0.5332 | 0.5234 |
| Sham_Vehicle | 7 day | 0.0030 | 0.0540 | 0.1665 | 0.0540 | 0.0700 | 0.3397 | 0.0116 | 0.0856 | 0.1515 | 0.2369 | 0.2115 | 0.7215 |
| Sham_Vehicle | 8 day | -0.0043 | 0.0520 | 0.1500 | 0.0520 | 0.1229 | 0.3609 | -0.0482 | 0.1180 | 0.1180 | 0.2328 | 0.2386 | 0.7991 |
| Sham_Vehicle | 9 day | -0.0142 | 0.0702 | 0.1514 | 0.0702 | 0.0713 | 0.3416 | 0.0196 | 0.0682 | 0.1016 | 0.3061 | 0.3497 | 0.6833 |

Supplemental table 4. Acute and chronic IR and radioprotectant treatment MEA metrics shown as mean and standard deviation (sd) for DoS normalized results.

| time | group1 | group2 | p.adj | p.adj.signif | MEA_metric |
| --- | --- | --- | --- | --- | --- |
| 1 day | Acute_amifostine | Acute_Vehicle | 0.00112 | ** | AE |
| 1 day | Acute_amifostine | Sham_Vehicle | 7.63e-05 | **** | AE |
| 1 day | Sham_amifostine | Sham_Vehicle | 0.00908 | ** | AE |
| 2 day | Acute_Vehicle | Sham_amifostine | 0.0116 | * | AE |
| 2 day | Acute_amifostine | Acute_Vehicle | 0.000432 | *** | AE |
| 2 day | Acute_amifostine | Sham_Vehicle | 6.43e-05 | **** | AE |
| 2 day | Sham_amifostine | Sham_Vehicle | 0.00236 | ** | AE |
| 3 day | Acute_Vehicle | Chronic_amifostine | 0.04 | * | AE |
| 3 day | Acute_Vehicle | Sham_amifostine | 0.00123 | ** | AE |
| 3 day | Acute_amifostine | Acute_Vehicle | 0.000175 | *** | AE |
| 3 day | Acute_amifostine | Chronic_Vehicle | 7.52e-08 | **** | AE |
| 3 day | Acute_amifostine | Sham_Vehicle | 1.85e-05 | **** | AE |
| 3 day | Chronic_Vehicle | Sham_amifostine | 1.54e-06 | **** | AE |
| 3 day | Chronic_amifostine | Chronic_Vehicle | 5.13e-05 | **** | AE |
| 3 day | Chronic_amifostine | Sham_Vehicle | 0.00492 | ** | AE |
| 3 day | Sham_amifostine | Sham_Vehicle | 0.000151 | *** | AE |
| 4 day | Acute_Vehicle | Sham_amifostine | 5.53e-05 | **** | AE |
| 4 day | Acute_amifostine | Acute_Vehicle | 8.59e-06 | **** | AE |
| 4 day | Acute_amifostine | Sham_Vehicle | 1.98e-06 | **** | AE |
| 4 day | Sham_amifostine | Sham_Vehicle | 1.53e-05 | **** | AE |
| 5 day | Acute_Vehicle | Sham_amifostine | 0.00095 | *** | AE |
| 5 day | Acute_amifostine | Acute_Vehicle | 8.89e-05 | **** | AE |
| 5 day | Acute_amifostine | Sham_Vehicle | 1.93e-05 | **** | AE |
| 5 day | Sham_amifostine | Sham_Vehicle | 0.00026 | *** | AE |
| 6 day | Acute_Vehicle | Sham_amifostine | 0.00902 | ** | AE |
| 6 day | Acute_amifostine | Acute_Vehicle | 0.000296 | *** | AE |
| 6 day | Acute_amifostine | Sham_Vehicle | 2.5e-05 | **** | AE |
| 6 day | Sham_amifostine | Sham_Vehicle | 0.00112 | ** | AE |
| 7 day | Acute_amifostine | Acute_Vehicle | 0.04 | * | AE |
| 7 day | Acute_amifostine | Chronic_Vehicle | 4.48e-05 | **** | AE |
| 7 day | Acute_amifostine | Sham_Vehicle | 0.00701 | ** | AE |
| 7 day | Chronic_Vehicle | Sham_amifostine | 0.000645 | *** | AE |
| 7 day | Chronic_amifostine | Chronic_Vehicle | 0.00161 | ** | AE |
| 7 day | Sham_amifostine | Sham_Vehicle | 0.0338 | * | AE |
| 8 day | Acute_amifostine | Chronic_Vehicle | 8.32e-05 | **** | AE |
| 8 day | Acute_amifostine | Sham_Vehicle | 0.0114 | * | AE |
| 8 day | Chronic_Vehicle | Sham_amifostine | 0.00289 | ** | AE |
| 8 day | Chronic_amifostine | Chronic_Vehicle | 0.00993 | ** | AE |
| 9 day | Acute_amifostine | Chronic_Vehicle | 0.000145 | *** | AE |
| 9 day | Acute_amifostine | Sham_Vehicle | 0.0277 | * | AE |
| 9 day | Chronic_Vehicle | Sham_amifostine | 0.0118 | * | AE |
| 9 day | Chronic_amifostine | Chronic_Vehicle | 0.0188 | * | AE |
| 3 day | Acute_amifostine | Chronic_Vehicle | 0.00212 | ** | AUNCC |
| 3 day | Acute_amifostine | Sham_Vehicle | 0.0203 | * | AUNCC |
| 4 day | Acute_amifostine | Sham_Vehicle | 0.0148 | * | AUNCC |
| 2 day | Acute_Vehicle | Sham_amifostine | 0.0364 | * | NBF |
| 3 day | Acute_amifostine | Sham_Vehicle | 0.0458 | * | NBF |
| 5 day | Acute_Vehicle | Sham_amifostine | 0.0314 | * | NBF |
| 5 day | Acute_amifostine | Acute_Vehicle | 0.0287 | * | NBF |
| 5 day | Acute_amifostine | Sham_Vehicle | 0.00152 | ** | NBF |
| 5 day | Sham_amifostine | Sham_Vehicle | 0.00243 | ** | NBF |
| 6 day | Acute_Vehicle | Sham_amifostine | 0.0127 | * | NBF |
| 6 day | Acute_amifostine | Acute_Vehicle | 0.0401 | * | NBF |
| 6 day | Acute_amifostine | Sham_Vehicle | 0.0128 | * | NBF |
| 6 day | Sham_amifostine | Sham_Vehicle | 0.00395 | ** | NBF |
| 2 day | Acute_Vehicle | Sham_amifostine | 0.0272 | * | NumB |
| 2 day | Acute_amifostine | Acute_Vehicle | 0.00847 | ** | NumB |
| 2 day | Acute_amifostine | Sham_Vehicle | 0.0076 | ** | NumB |
| 2 day | Sham_amifostine | Sham_Vehicle | 0.0272 | * | NumB |
| 3 day | Acute_Vehicle | Chronic_amifostine | 0.00973 | ** | NumB |
| 3 day | Acute_Vehicle | Sham_amifostine | 0.000506 | *** | NumB |
| 3 day | Acute_amifostine | Acute_Vehicle | 0.000518 | *** | NumB |
| 3 day | Acute_amifostine | Chronic_Vehicle | 4.05e-06 | **** | NumB |
| 3 day | Acute_amifostine | Sham_Vehicle | 0.000381 | *** | NumB |
| 3 day | Chronic_Vehicle | Sham_amifostine | 4.73e-06 | **** | NumB |
| 3 day | Chronic_amifostine | Chronic_Vehicle | 0.000101 | *** | NumB |
| 3 day | Chronic_amifostine | Sham_Vehicle | 0.00683 | ** | NumB |
| 3 day | Sham_amifostine | Sham_Vehicle | 0.000374 | *** | NumB |
| 4 day | Acute_Vehicle | Sham_amifostine | 0.00082 | *** | NumB |
| 4 day | Acute_amifostine | Acute_Vehicle | 0.00092 | *** | NumB |
| 4 day | Acute_amifostine | Sham_Vehicle | 0.00093 | *** | NumB |
| 4 day | Sham_amifostine | Sham_Vehicle | 0.00092 | *** | NumB |
| 5 day | Acute_Vehicle | Sham_amifostine | 0.00637 | ** | NumB |
| 5 day | Acute_amifostine | Acute_Vehicle | 0.01 | * | NumB |
| 5 day | Acute_amifostine | Sham_Vehicle | 0.01 | * | NumB |
| 5 day | Sham_amifostine | Sham_Vehicle | 0.00637 | ** | NumB |
| 6 day | Acute_Vehicle | Sham_amifostine | 0.047 | * | NumB |
| 6 day | Acute_amifostine | Sham_Vehicle | 0.0323 | * | NumB |
| 6 day | Sham_amifostine | Sham_Vehicle | 0.0253 | * | NumB |
| 7 day | Acute_amifostine | Chronic_Vehicle | 0.0461 | * | NumB |
| 7 day | Chronic_Vehicle | Sham_amifostine | 0.01 | * | NumB |
| 7 day | Chronic_amifostine | Chronic_Vehicle | 0.0476 | * | NumB |
| 8 day | Chronic_Vehicle | Sham_amifostine | 0.0189 | * | NumB |
| 1 day | Acute_amifostine | Sham_Vehicle | 0.000236 | *** | WMFR |
| 1 day | Sham_amifostine | Sham_Vehicle | 0.00927 | ** | WMFR |
| 2 day | Acute_amifostine | Acute_Vehicle | 0.00571 | ** | WMFR |
| 2 day | Acute_amifostine | Sham_Vehicle | 0.000497 | *** | WMFR |
| 2 day | Sham_amifostine | Sham_Vehicle | 0.0108 | * | WMFR |
| 3 day | Acute_Vehicle | Sham_amifostine | 0.00452 | ** | WMFR |
| 3 day | Acute_amifostine | Acute_Vehicle | 0.0015 | ** | WMFR |
| 3 day | Acute_amifostine | Chronic_Vehicle | 8.29e-07 | **** | WMFR |
| 3 day | Acute_amifostine | Sham_Vehicle | 8.4e-05 | **** | WMFR |
| 3 day | Chronic_Vehicle | Sham_amifostine | 6.2e-06 | **** | WMFR |
| 3 day | Chronic_amifostine | Chronic_Vehicle | 9.59e-05 | **** | WMFR |
| 3 day | Chronic_amifostine | Sham_Vehicle | 0.00489 | ** | WMFR |
| 3 day | Sham_amifostine | Sham_Vehicle | 0.000302 | *** | WMFR |
| 4 day | Acute_Vehicle | Sham_amifostine | 0.00267 | ** | WMFR |
| 4 day | Acute_amifostine | Acute_Vehicle | 0.000441 | *** | WMFR |
| 4 day | Acute_amifostine | Sham_Vehicle | 0.00011 | *** | WMFR |
| 4 day | Sham_amifostine | Sham_Vehicle | 0.000849 | *** | WMFR |
| 5 day | Acute_Vehicle | Sham_amifostine | 0.00443 | ** | WMFR |
| 5 day | Acute_amifostine | Acute_Vehicle | 0.00119 | ** | WMFR |
| 5 day | Acute_amifostine | Sham_Vehicle | 0.000235 | *** | WMFR |
| 5 day | Sham_amifostine | Sham_Vehicle | 0.00116 | ** | WMFR |
| 6 day | Acute_amifostine | Acute_Vehicle | 0.00539 | ** | WMFR |
| 6 day | Acute_amifostine | Sham_Vehicle | 5e-04 | *** | WMFR |
| 6 day | Sham_amifostine | Sham_Vehicle | 0.00794 | ** | WMFR |
| 7 day | Acute_amifostine | Chronic_Vehicle | 0.000382 | *** | WMFR |
| 7 day | Acute_amifostine | Sham_Vehicle | 0.0329 | * | WMFR |
| 7 day | Chronic_Vehicle | Sham_amifostine | 0.00171 | ** | WMFR |
| 7 day | Chronic_amifostine | Chronic_Vehicle | 0.00182 | ** | WMFR |
| 8 day | Acute_amifostine | Chronic_Vehicle | 0.000374 | *** | WMFR |
| 8 day | Acute_amifostine | Sham_Vehicle | 0.0362 | * | WMFR |
| 8 day | Chronic_Vehicle | Sham_amifostine | 0.016 | * | WMFR |
| 8 day | Chronic_amifostine | Chronic_Vehicle | 0.0106 | * | WMFR |
| 9 day | Acute_amifostine | Chronic_Vehicle | 0.000201 | *** | WMFR |
| 9 day | Acute_amifostine | Sham_Vehicle | 0.0272 | * | WMFR |
| 9 day | Chronic_Vehicle | Sham_amifostine | 0.0199 | * | WMFR |
| 9 day | Chronic_amifostine | Chronic_Vehicle | 0.023 | * | WMFR |

Supplemental table 5. All significant IR and radioprotectant pairwise comparisons by Holm-Bonferroni test.
